# The effects of mutations on gene expression and alternative splicing: a case study of EMS-induced heritable mutations in the microcrustacean *Daphnia*

**DOI:** 10.1101/2023.02.28.530525

**Authors:** Marelize Snyman, Sen Xu

**Author notes:** Correspondence author. 501 S Nedderman Dr., Arlington, Texas, USA 76019.

## Abstract

Understanding the relationship between mutations and their genomic and phenotypic consequences has been a longstanding goal of evolutionary biology. However, few studies have investigated the impact of mutations on gene expression and alternative splicing on the genome-wide scale. In this study, we aim to bridge this knowledge gap by utilizing whole-genome sequencing data and RNA sequencing data from 16 OP *Daphnia* mutant lines to investigate the effects of EMS-induced mutations on gene expression and alternative splicing. Using rigorous analyses of mutations, expression changes, and alternative splicing, we show that trans-effects are the major contributor to the variance in gene expression and alternative splicing between the wildtype and mutant lines, whereas cis mutations only affected a limited number of genes and do not always alter gene expression. Moreover, we show that there is a significant association between DE genes and exonic mutations, indicating that exonic mutations are an important driver of altered gene expression.

## Introduction

One of the fundamental goals of evolutionary biology is to understand the genomic and phenotypic consequences of mutations. As mutations are rare, long-term mutation accumulation (MA) experiments in a variety of different model organisms, e.g. *Drosophila* (Keightley *et al*. 2009, Krasovec 2021), *Caenorhabditis* (Denver *et al*. 2009), *Arabidopsis* (Ossowski *et al*. 2010) and *Daphnia* (Bull *et al*. 2019, Flynn *et al*. 2017, Omilian *et al*. 2006, Xu *et al*. 2012) have been instrumental in our understanding of mutational consequences. These MA experiments minimized the power of selection on mutations and allow for the persistence of non-lethal mutations in a nearly unbiased manner, offering insight into the genome-wide rates, spectrum, and fitness effects of spontaneous mutations. The findings of these studies have yielded important implications for our understanding of diseases, the origin of genetic diversity, and molecular evolution in general.

To date, many MA studies have moved beyond simply measuring the mutation rate, spectrum, and fitness effects on the organismal level. For example, MA studies have been utilized to study transposable element mutation rates (Díaz-González and Domínguez 2020, Ho *et al*. 2021), enzymatic activity (Aquadro *et al*. 1990, Harada 1995), methylation frequency (Denkena *et al*. 2021, Jiang *et al*. 2014), phenotypic plasticity (Latta *et al*. 2015), and patterns of genotype-environment interactions (Chu and Zhang 2021, Scheffer *et al*. 2022, Xu 2004).

However, few studies have examined the impacts of mutations on gene expression, which could have dramatic phenotypic consequences (see below). Furthermore, while most MA experiments focus on spontaneous mutations, few studies have paid attention to environmentally induced mutations. As anthropogenic stressors such as chemical pollutants (e.g., Chernobyl disaster in 1986, Fukushima nuclear disaster in 2011) are impacting various ecosystems at a dramatically increased rate since the start of the Industrial Revolution (Landrigan *et al*. 2018), it has become imperative to understand the genetic consequences of these mutations induced by environmental agents such as chemical mutagens. In this study, we aim to bridge this knowledge gap by investigating the impact that ethyl-methanesulfonate (EMS)-induced inheritable mutations exert on genome-wide gene expression and alternative splicing.

Gene expression is a major link between an organism’s genotype and phenotype. Numerous studies have demonstrated the critical role of altered gene expression in phenotypic variation such as the beak morphology in Darwinian finches (Abzhanov *et al*. 2006), cold adaptation in lizards (Josephs 2021), melanin production in *Drosophila* (Rebeiz *et al*. 2009), body mass due to latitudinal adaptation and color pattern in mice (Mack *et al*. 2018, Manceau *et al*. 2011), and adaptation to predation in Trinidadian guppies (Ghalambor *et al*. 2015).

Variation in gene expression can arise through cis and trans mutations. Cis mutations are often located within a gene, or up- and downstream of a gene, affecting the interaction between the gene and its regulators (e.g., promotors and enhancers). On the other hand, trans mutations can be located anywhere within the genome (other than within or nearby the gene of interest) and mediate the expression of genes through the use of e.g., transcription factors and long noncoding RNAs (Albert *et al*. 2018, Lewis *et al*. 2014, Vande Zande *et al*. 2022). Trans mutations are thought to be primarily responsible for the variation in gene expression within species since trans factors represent a much larger mutational target and are thus more likely to arise than cis mutations (Chen *et al*. 2015, Coolon *et al*. 2014, Gruber *et al*. 2012, Metzger *et al*. 2016, Rhoné *et al*. 2017, Wittkopp *et al*. 2004). However, due to the pleiotropic effects of trans mutations on expression (e.g., affecting multiple genes at the same time), trans mutations are often selected against as populations adapt and diverge. With cis mutations mostly affecting the expression of specific genes, they survive selection more often than trans mutations. Therefore, cis mutations rather than trans mutations are mostly found to be responsible for gene variation between populations and between species (Coolon *et al*. 2015, Emerson *et al*. 2010, Schaefke *et al*. 2013).

These theories and empirical observations in natural populations lead us to expect that de novo mutations (spontaneous or environmentally induced) upon arising in the genome should exhibit stronger trans effects than cis effects. However, this hypothesis remains largely untested due to previous work focusing on utilizing targeted mutagenesis to examine the effects of cis mutations in a variety of different organisms (Hornung *et al*. 2012, Kwasnieski *et al*. 2012, Maricque *et al*. 2017, Melnikov *et al*. 2012, Metzger *et al*. 2015, Patwardhan *et al*. 2009, Sharon *et al*. 2012). We suggest that utilizing chemical mutagenesis throughout the genome provides an important alternative approach to understanding how cis and trans mutations affect gene expression on the genome-wide level.

Furthermore, a very much ignored aspect of mutations’ impact on gene expression resides in alternative splicing. Alternative splicing is critically involved in the origin of phenotypes, e.g., the flowering time of *Arabidopsis* (Macknight *et al*. 2002), seed development in sunflowers (Smith *et al*. 2018), pigmentation of cichlid fishes (Terai *et al*. 2003), and thermogenesis in mice (Vernia *et al*. 2016). With the involvement of trans-acting splicing factors and cis-acting regulatory motifs, alternative splicing is often highly regulated and vulnerable to mutations. One can imagine that point mutations can occur in both introns and exons, leading to the alteration of existing splice sites or the generation of new ones, and affecting splicing enhancers and silencers (Anna and Monika, 2018). However, it remains unclear to what extent alternative splicing is affected by trans mutations.

To examine the effects of mutations on gene expression and alternative splicing on a genome-wide scale, we employed the chemical mutagen ethyl methanesulfonate (EMS) to induce heritable mutations in the microcrustacean *Daphnia*. EMS induces DNA damage through the alkylation of guanine, causing O^6^ ethylguanine to mispair with thymine instead of cytosine in subsequent replications, resulting in EMS-induced mutations mainly consisting of G:C to A:T transitions (Greene *et al*. 2003). EMS-induced mutations are randomly distributed throughout the genome, enabling the generation of loss or gain of function mutants and weak nonlethal alleles (Greene *et al*. 2003, Lee *et al*. 2003).

We have established an EMS mutagenesis protocol for *Daphnia* (Snyman *et al*. 2021), where we showed that by exposing *Daphnia* to EMS, the base substitution rate reached 1.17×10^- 6^ and 1.75 ×10^-6^ per base per generation for 10mM and 25mM EMS concentrations, respectively (Snyman *et al*. 2021). These EMS-induced mutation rates are greatly elevated compared to the spontaneous base substitution rate estimates in *Daphnia* of 2.30×10^−9^ and 7.17 × 10^-9^ per base per generation (Flynn *et al*. 2017, Keith *et al*. 2016). The EMS-induced mutations were also randomly distributed in different types of genomic regions (e.g., exon, intron, intergenic regions).

In this study, we selected an obligately parthenogenetic *Daphnia* isolate to examine the impact of EMS-induced heritable mutations on gene expression and alternative splicing. A major motivation for using an obligately parthenogenetic isolate is that we can avoid the maternal effects imposed by exposure to EMS in the ancestral mutant as much as possible by examining the asexual offspring of later generations that are genetically identical (albeit extremely rare spontaneous mutations) to the ancestors. *Daphnia* typically reproduces by cyclical parthenogenesis. Under favorable environmental conditions such as high food abundance and low population density, *Daphnia* females asexually produce diploid subitaneous eggs, giving rise to genetically identical daughters **(Figure 1)**. Under stressful environmental conditions (e.g., high population density, lack of food availability), females switch to sexual reproduction, producing haploid eggs. Males also emerge in populations under environmental stress (hatched from diploid subitaneous eggs), as sex is environmentally determined in *Daphnia* (Gorr *et al*. 2006, Tatarazako *et al*. 2003). Sexual reproduction generates dormant embryos deposited into a protective case (i.e., ephippium), which can resume development once the environment becomes favorable (Frisch *et al*. 2014). However, for obligately parthenogenetic *Daphnia*, females under stress can asexually produce diploid resting eggs (Xu *et al*. 2022).

**Figure 1.**
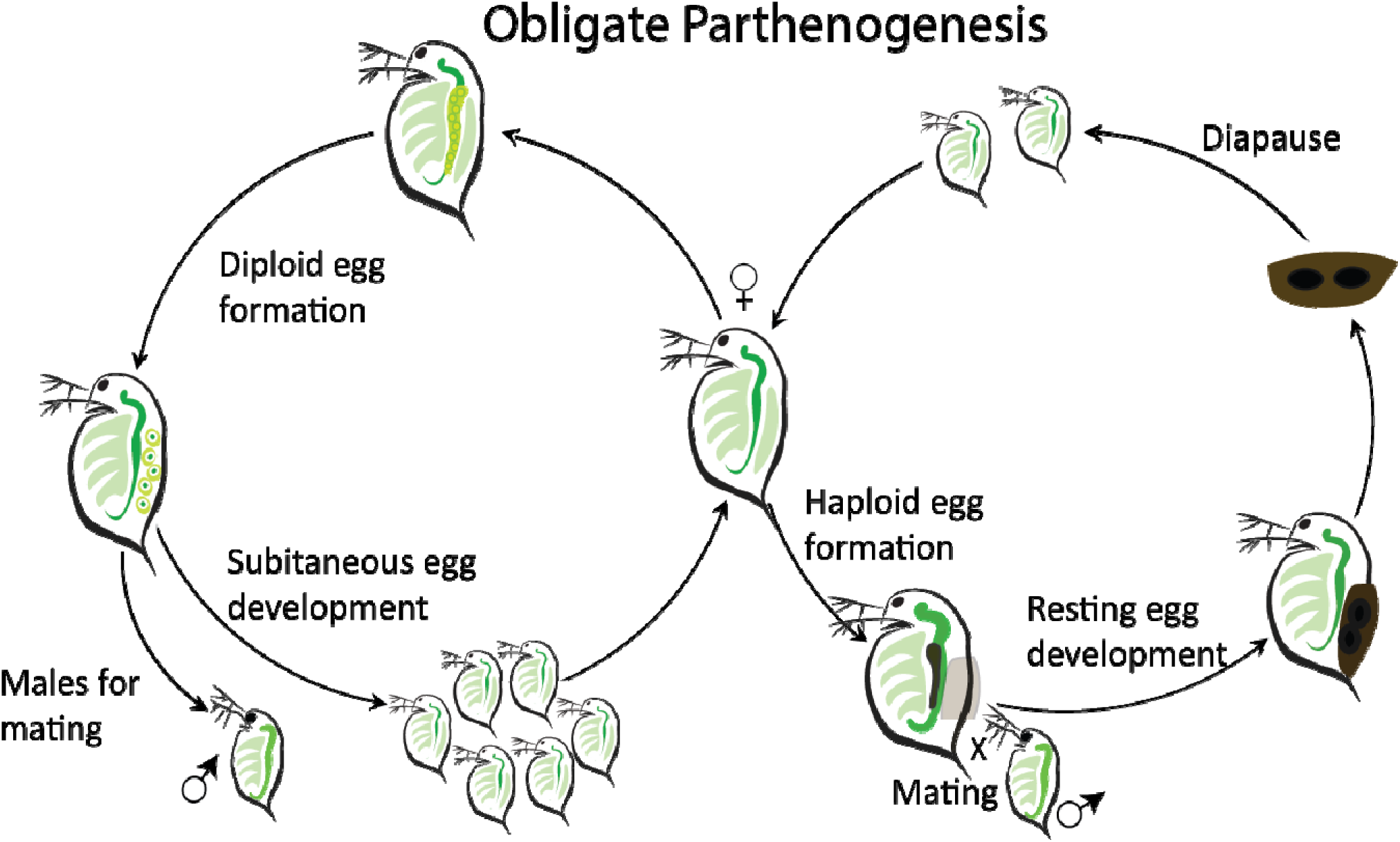
The life cycle of obligately parthenogenetic *Daphnia*.

In this study, we examined the gene expression and alternative splicing mechanisms functional in 16 *Daphnia* mutant lines derived from the same ancestor in comparison to the wildtype using whole-genome DNA and RNA sequencing data. Our main goals were three-fold. First, we investigated the alteration of gene expression (DS) and alternative splicing in the mutant lines due to the impact of EMS-induced mutations. Second, we examined the contribution of cis and trans mutations on differential gene expression and alternative splicing. Third, we were interested in whether any genomic regions carrying cis mutations were significantly associated with DE and AS.

## Materials and Methods

### Sampling and maintenance of isolates

An obligately parthenogenetic *Daphnia* isolate collected from a pond in the US was used in this study. This isolate has been kept as a clonally reproducing line in artificial lake (Kilham *et al*. 1998) water under a 16:8 (light: dark) cycle at 18°C and fed with the green algae, *Scenedesmus obliquus* twice a week.

### Generating Daphnia mutants using EMS

To generate *Daphnia* mutants with heritable mutations from the wildtype, we used EMS to induce germline mutations following the established methodology in *Daphnia* (Snyman *et al*. 2021). Specifically, eight sexually mature *Daphnia* females (at the asexually reproducing stage for subitaneous eggs) were exposed to 25 mM ethyl methanesulfonate for 4 hours to introduce mutations into their oocytes. After exposure, these females were individually isolated **(Figure 2)**. We collected two progenies (G_1_s) in the first asexual brood from each of the eight exposed females because each G_1_ was derived from an oocyte that was individually mutagenized, thus all the G_1_s are genetically distinct. In total, we established 16 mutant lines.

**Figure 2.**
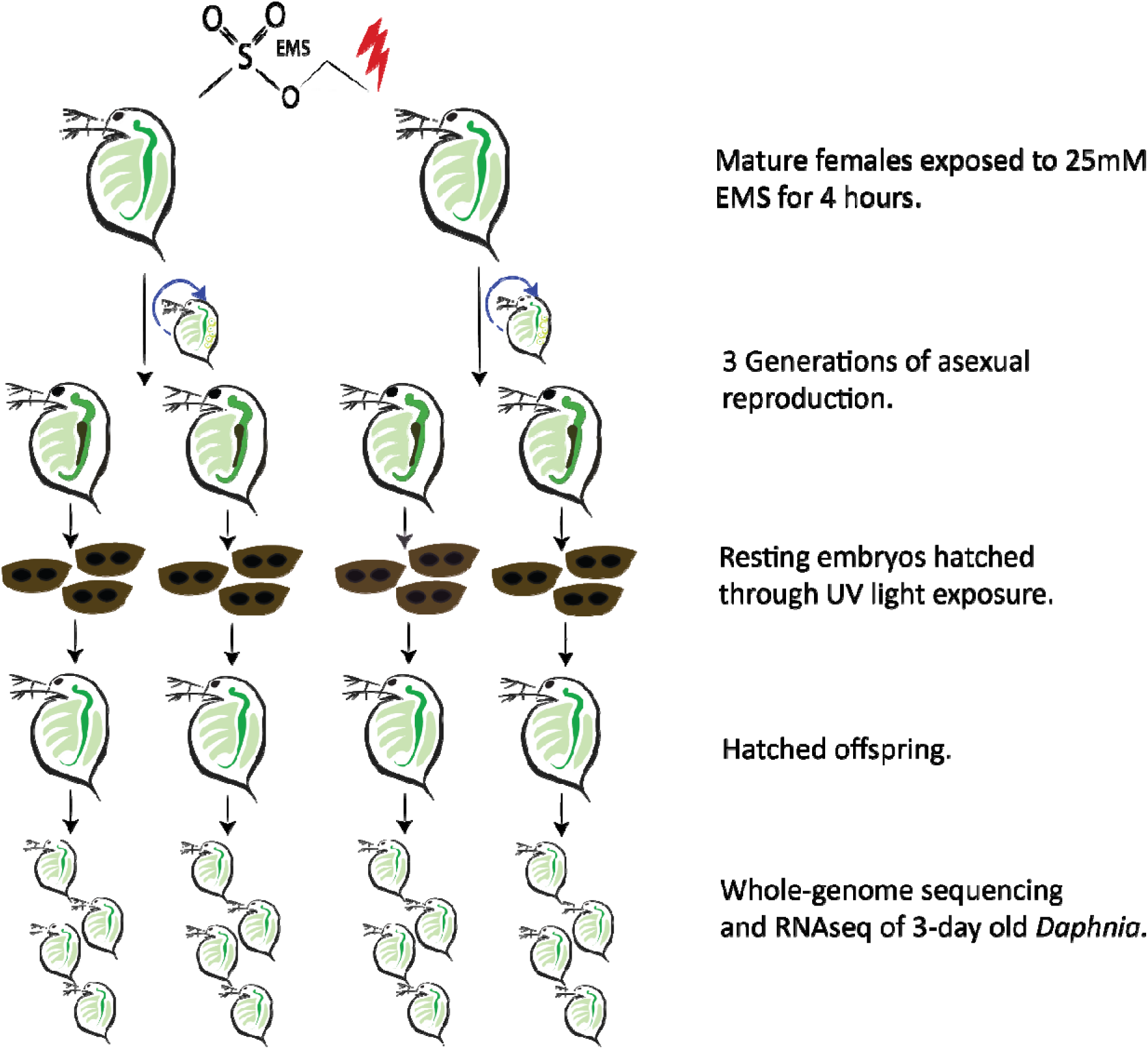
Illustration of exposure method to generate EMS mutant lines.

To avoid maternal effect (i.e., from exposure to EMS) on gene expression in the offspring, each of the mutant lines was propagated asexually for at least 3 generations to reach a high enough animal density to induce the parthenogenetic production of resting eggs. 5-10 ephippia (each containing 2 resting eggs) were collected from each mutant line, and the decapsulated resting eggs were hatched using the protocol established by Luu *et al*. 2020.

Briefly, the collected resting embryos were kept in the dark at 18°C for two weeks before being placed under UV light to stimulate embryo development. If no sign of embryo development was observed after five days, the resting embryos were placed back in the dark for 2 weeks and then re-exposed to UV light until at least one resting embryo hatched per mutant line. The hatched mutant offspring was then asexually propagated until a high enough animal density was reached for DNA and RNA extraction.

### DNA extraction and whole-genome DNA sequencing

A total of 30-40 clonal offspring were collected for each mutant line for DNA extraction using a CTAB (Cetyl Trimethyl Ammonium Bromide) method (Doyle and Doyle 1987). DNA quality and concentration were assessed by electrophoresis on a 2% agarose gel and a Qubit 4.0 Fluorometer (Thermo Scientific, Waltham, MA, USA). DNA sequencing libraries were prepared following standard MGI sequencing library protocol by the Beijing Genomics Institute (BGI, Cambridge, MA, USA). All 16 mutant lines were sequenced on an MGI DNBseq platform with 150-bp paired-end reads, with a targeted sequencing coverage of 30X per line. The raw DNA sequence data were deposited at NCBI SRA under PRJNA892919.

### RNA extraction and transcriptomic sequencing

Experimental animals were maintained in the same environmental conditions i.e., 18L with a 16:8 light/dark cycle, and three replicates of 2 or 3-day old offspring were collected from the wild-type (i.e., control), and each mutant line. RNA was extracted using the Promega SV Total RNA Isolation kit (Madison, WI, USA) following the manufacturer’s instructions. RNA quality was examined by electrophoresis on a 2% agarose gel, and RNA concentration was measured using a Qubit 4.0 Fluorometer (Thermo Scientific, Waltham, MA, USA). RNA sequencing libraries were prepared using the NEBNext Ultra II RNA Library Prep Kit for Illumina (Ipswich, MA, USA). Transcriptomic sequencing was done by Novogene Corporation Inc. (Sacramento, CA, USA) following standard Illumina sequencing protocol. Each library was sequenced on an Illumina NovaSeq6000 platform with at least 20 million 150-bp paired-end reads. The raw RNA sequence data were deposited at NCBI SRA underPRJNA892982.

### Quality control and mapping

Quality of the raw reads was examined using FastQC (Andrews 2010). No adapter contamination was observed for the whole-genome sequencing data, thus further analysis was completed using the raw reads. Our RNAseq dataset showed adapter contamination. To address this issue Trimmomatic v.0.39 (Bolger *et al*. 2014) was used to perform adapter trimming and quality filtering using the following parameters: ILLUMINACLIP adapter.fasta:2:30:10:2:keepBothReads LEADING:3 TRAILING:3 MINLEN:36. The following Illumina adapter sequences were trimmed, read 1: AGATCGGAAGAGCACACGTCTGAACTCCAGTCA, read 2: AGATCGGAAGAGCGTCGTGTAGGGAAAGAGTGT. Lastly, reads were reassessed using FastQC to confirm the removal of low-quality reads and adapter sequences.

### Identification of EMS-induced heritable mutations

We identified heterozygous EMS-induced heritable mutations following the EMS mutagenesis analysis protocol established by Snyman *et al*. (2021). Since these mutant lines are obligate parthenogens (i.e., no segregation and no sexual reproduction) all EMS-induced mutations should be in the heterozygous state, ignoring rare ameiotic gene conversions (Omilian et al 2006). We used the Burrows-Wheeler Alignment Tool BWA-MEM version 0.7.17 (Li and Durbin 2009) with default parameters to align the whole-genome DNA sequencing raw reads of each mutant line to the *Daphnia pulicaria* reference genome (Jackson *et al*. 2021). SAMtools (Li *et al*. 2009) was used to remove reads mapped to multiple locations, and the MarkDuplicates function of Picard tools (http://broadinstitute.github.io/picard/) was used to locate and tag PCR duplicates. We used BCFtools (Li 2011) mpileup and call functions with default parameters to generate genotype likelihoods and genotype calls in a VCF file. The following additional FORMAT and INFO tags were added to the VCF file: AD (allelic depth), DP (number of high-quality bases), ADF (allelic depth on forward strand), and ADR (allelic depth on reverse strand). Tentative EMS-induced mutations for all treatment lines were further filtered with BCFtools’ filter function to retain only single nucleotide polymorphisms (SNPs) with a sequencing depth (DP) >= 10 and <= 60, quality score (QUAL) >= 20, and a distance of at least 50-bp from an indel.

Furthermore, a custom python script (https://github.com/Marelize007/Functional_impact_of_EMS-induced_mutations) was used to filter the tentative mutations. As described by Snyman *et al*. (2021), the final set of mutations was identified using a consensus method. Briefly, all genotype data from the 16 EMS mutant lines were added to one VCF file, and a consensus genotype was established by majority rule. Out of the 16 mutant lines, at least 12 mutant lines had to agree to generate a consensus genotype call per site. This allowed for the inclusion of mutation sites where up to 3 out of the 16 mutant lines had no genotype call due to inadequate depth. A tentative mutation was identified if the mutant line showed a different genotype than the consensus. All final mutations had to be supported by at least two forward and two reverse reads to limit false positives due to allele drop and inadequate sequence coverage.

### Differential expression analysis

The trimmed RNAseq reads were mapped to the *D. pulicaria* (Jackson *et al*. 2021) reference genome utilizing the STAR aligner (Dobin *et al*. 2013) with default parameters. Reads mapped to multiple locations in the genome were removed using SAMtools (Li *et al*. 2009), and raw transcript counts were obtained for each sample using featureCounts (Liao *et al*. 2014).

Differential expression analysis was performed using DESeq2 v.1.34.0 (Love *et al*. 2014). First, the regularized log (rlog) transformation function in DEseq2 was used to normalize the mapped read counts. Next, differentially expressed genes were determined for each mutant line using the Wald negative binomial test with the design formula ∼ genotype, where genotype represents either the mutant or wild-type genotype. The Benjamini-Hochberg method (FDR < 0.05) was used to adjust the p-values for multiple testing, and genes with a p-value <0.05 were considered significantly differentially expressed. Differentially expressed genes were additionally filtered according to fold change. Genes with a fold-change > 1.5 were considered upregulated, and < - 1.5 were considered downregulated in the mutant lines compared to the wildtype (i.e., control).

### Mutation rate calculation

The formula, *µ = m/n *l* was used to calculate the per site per generation mutation rate for all 16 mutant lines, where *m* is the total number of mutations identified in each line, *n* is the total number of genomic sites with a sequencing depth >=10, and <=60, QUAL >= 20, and where each site is at least 50-bp from the nearest indel in each mutant line. Furthermore, *l* represents the number of generations. To calculate the per gene per generation mutation rate, we used the formula *µ_g_ = m_g_/n_g_ *1,* where *m_g_* represents the total number of mutations detected in genic regions (including UTRs, exons, and introns), *n_g_* represents the total number of genes analyzed in each mutant line, and *l* represents one generation. The non-synonymous mutation rate was calculated utilizing the same formula, except *m_g_* represented the number of non-synonymous mutations per mutant line.

### Annotation of EM-induced mutations

Functional annotation based on genomic locations and effect prediction of EMS-induced heritable mutations were done using the cancer mode (-cancer) of SnpEff version 4.0 with default parameters (Cingolani *et al*. 2012). This mode was utilized since it allowed direct comparison between the EMS mutant genotypes and the wild type.

### Alternative splicing analysis

We used the tool rMATS v4.1.1 (Shen *et al*. 2014) to detect alternatively spliced (AS) events using reads mapped to both exons and splice junctions. The following alternatively spliced events were detected: skipped exon (SE), alternative 5’ splice site (A5SS), alternative 3’ splice site (A3SS), retained intron (RI), and mutually exclusive exons (MXE). SE events take place when an exon along with its flanking introns are spliced out, and A3SS and A5SS result when different parts of exons are either included or excluded from the resulting transcript. During RI events, introns are retained; during MXE events, only one of two exons are retained in the resulting mRNA (Pohl *et al*. 2013, Wang *et al*. 2015). Alternatively spliced events had to be supported by at least four uniquely mapped reads and have a minimum anchor length of 10nt. Additionally, the Benjamini-Hochberg adjusted (FDR < 0.05) p-value had to be less than 0.05, and the difference in exon inclusion level (Δ|ψ|) greater than 5% (Shen *et al*. 2014, Suresh *et al*. 2020).

## Results

### Whole-genome DNA and RNA sequencing data

A total of 16 EMS-induced mutant lines were whole-genome sequenced with 150-bp Illumina paired-end reads **(Supplementary Table S1)**. We obtained ∼6GB of raw DNA sequence data for each mutant line, containing an average of ∼34 million reads (SD = 3.7) per sample. Read trimming was not performed since the results from the FastQC analysis did not reveal any adapter contamination. After removing PCR duplicates and reads mapped to multiple locations, each mutant line had on average ∼29 million (SD = 2.5) mapped reads with an average coverage of 25 reads (SD = 2) per site.

Furthermore, RNA sequencing data were obtained for three replicates per mutant line and one wildtype (control) line, totaling 51 samples. Approximately 20 million (SD = 4.9 million) 150-bp paired-end reads were obtained per sample. After adapter trimming and quality control, ∼99% of the reads were kept and mapped to the *D. pulicaria* reference genome (Jackson *et al*. 2021) with an average mapping rate of 95% (SD = 2.37%, **Supplementary Table S2)**.

### EMS-induced heritable mutation rate and spectrum

To identify EMS-induced mutations in these mutant lines, we used a rigorous bioinformatic EMS mutation identification pipeline that a previous study tested with PCR mutation verification (false positive rate < 0.05, Snyman *et al*. 2021). The number of EMS-induced heritable heterozygous mutations in each mutant line ranged from 125 to 576 (median =152), translating to a base substitution rate ranging from 1.51 to 6.98×10^-6^ (median= 1.84×10^-6^) per site per generation. The number of mutations found within genic regions (including UTRs, exons, and introns) ranged from 49 to 320 (median = 74) in each line, resulting in a mutation rate of 2.0×10^- 3^ to 1.3×10^-2^ (median =3.1×10^-3^) per gene per generation **(Figures 3A and 3B, Supplementary Table S3)**.

**Figure 3.**
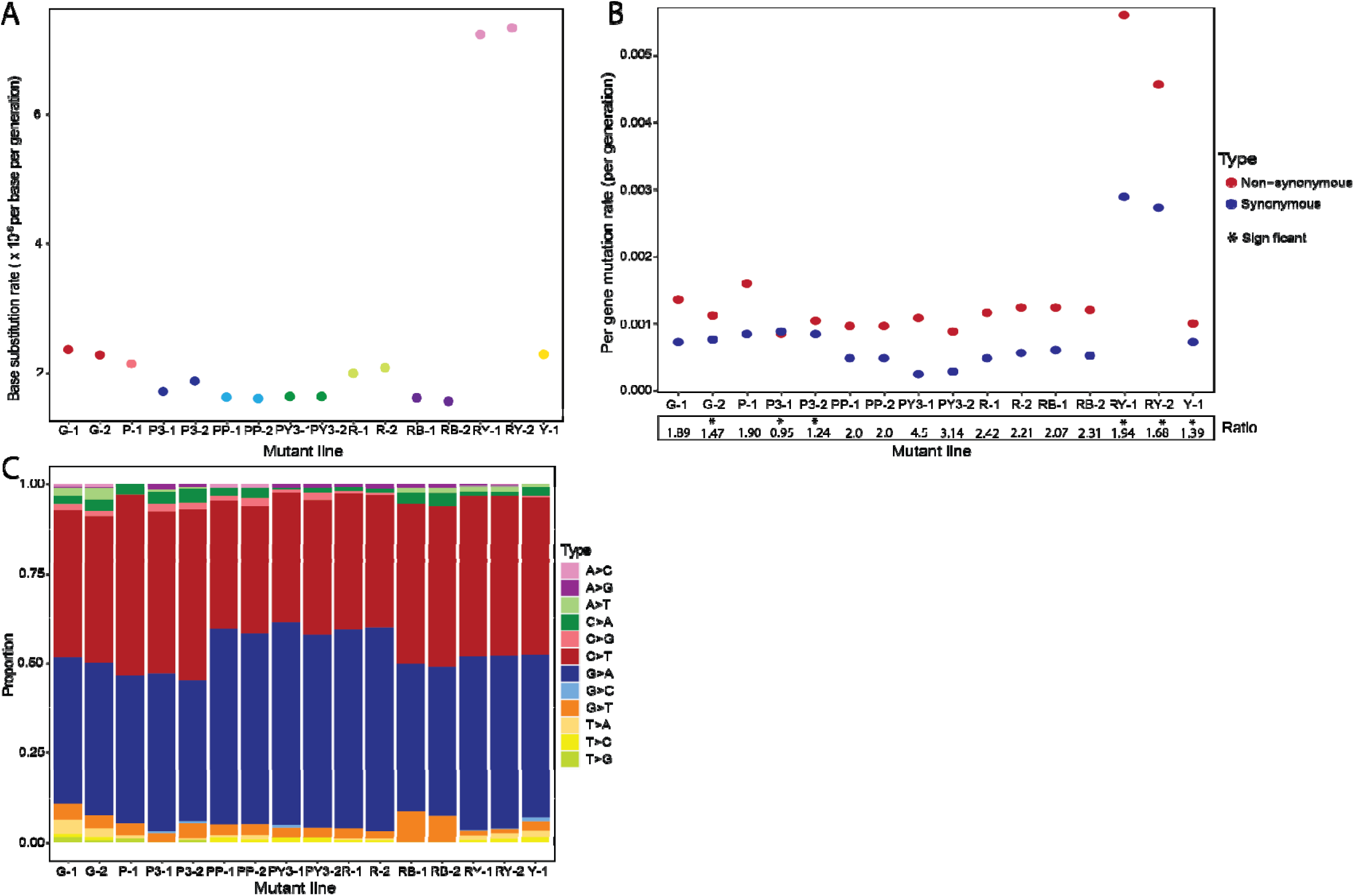
(A) The base substitution rates. (B) Non-synonymous and synonymous mutation rate, and non-synonymous *vs* synonymous mutation ratio in all mutant lines. (C) The proportion of base substitutions caused by EMS-induced mutations.

Across all 16 EMS-induced mutant lines the number of non-synonymous mutations ranged from 21 to 140 (median = 29), resulting in a median non-synonymous mutation rate of 1.14 ×10^-3^ per gene per generation **(Figure 3B)**. The number of synonymous mutations ranged from 6 to 72 (median = 17, **Supplementary Table S3**) across mutant lines. Assuming a random distribution of EMS-induced mutations, we expect to see a non-synonymous *vs* synonymous mutation ratio of 3:1. However, in 6 of the mutant lines we observed a significant deviation (one proportion Z test p-value < 0.05) from the expected 3:1 ratio (**Figure 3 B**, Graur and Li 2000).

Lastly, in accordance with previous work on *Daphnia* (Snyman *et al*. 2021) and other model organisms such as *C. elegans* (Flibotte *et al*. 2010) and *D. melanogaster* (Pastink *et al*. 1991), the majority of EMS-induced mutations across all 16 mutant lines were G:C to A:T transitions (mean = 90%, SD = 4%), yielding a transition-transversion ratio greater than 5.06 for all mutant lines **(Figure 3C, Supplementary Table S4)**.

### Differentially expressed genes in mutant lines

To assess how EMS-induced mutations altered the transcript abundance across the genome, we examined the transcriptomic dataset derived from 16 mutant lines in comparison to the wildtype. Our principal component analysis based on the normalized read counts (rlog transformation) showed the tight clustering of replicates derived from the same mutant, suggesting strong overall similarity among replicates of the same mutant background (**Figure 4A)**. The top two principal components (PC) were together responsible for >50% of the variance, with PC1 accounting for 32% and PC2 for 21%. It was not clear what biological factors these two principal components potentially represent. Most likely, they jointly captured the variability between mutants, between sibling mutant lines, and between replicates.

**Figure 4.**
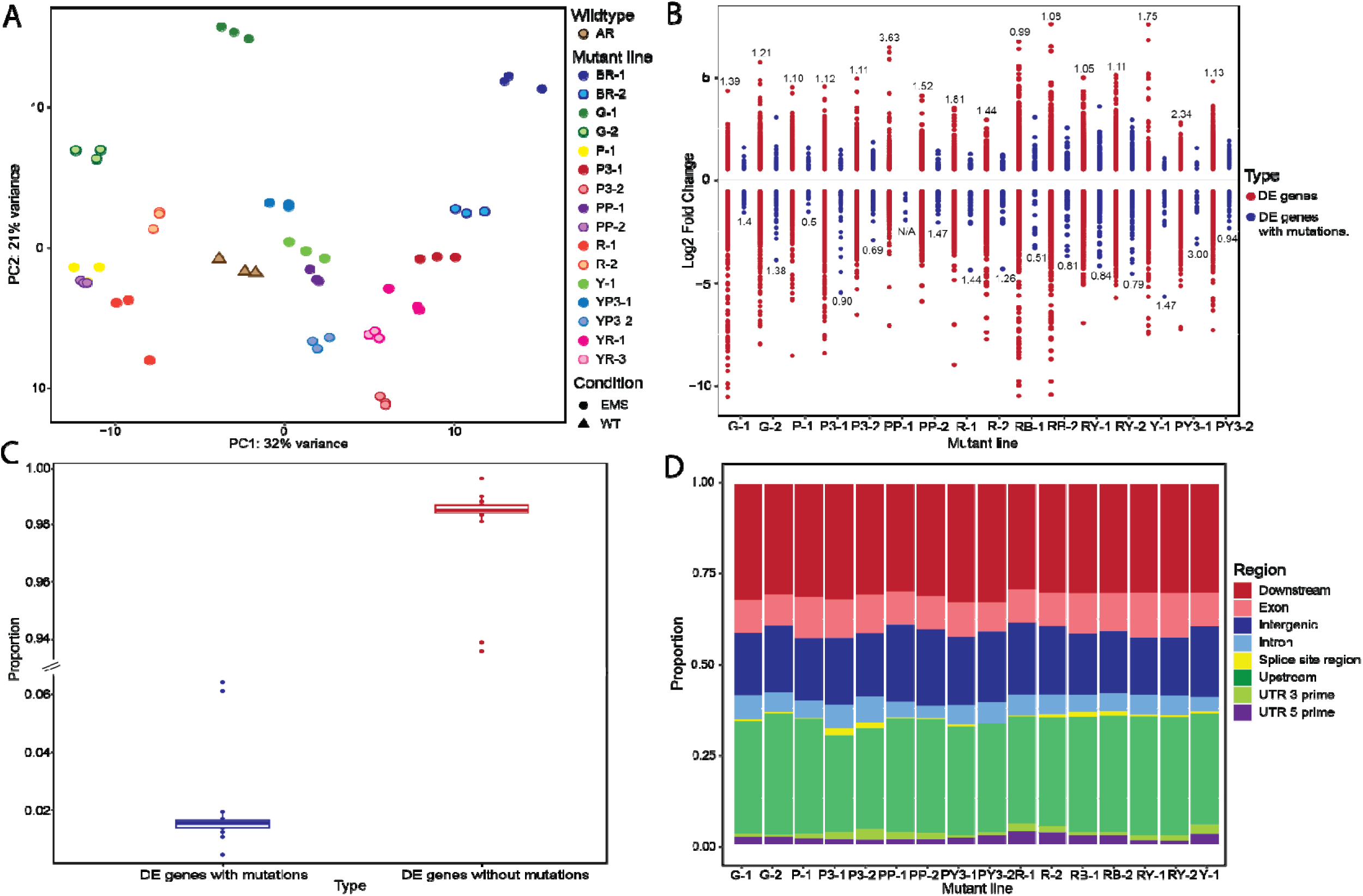
(A) PCA plot based on transcriptomic data of mutant lines. (B) Log2 fold change of differentially expressed (DE) genes (red dots) and DE genes with mutations (blue dots) with the ratio of down- *vs.* upregulated DE genes for each line. (C) Boxplot illustrating the proportions of differentially expressed (DE) genes with and without mutations. (D) Distribution of EMS-induced mutations in different genomic regions.

Our differential expression analysis contrasting each mutant line against the wildtype revealed that the total number of differentially expressed (DE) genes ranged from 1176 to 6606 across all lines (p < 0.05 with FDR < 0.05), with a median of 3545. In nearly all mutant lines the ratio of down- *vs.* upregulated genes was greater than 1 (1.05-3.63, **Figure 4B, Supplementary Table S5)**, demonstrating that downregulated genes were more abundant than upregulated genes. Only one mutant line (RB1) showed a slight majority of upregulated genes with a ratio of 0.99.

Next, we assessed the number of DE genes carrying an EMS-induced mutation as well as DE genes free of a mutation. We defined that a gene carries an EMS-induced mutation if a mutation was located 5kb upstream or downstream of the gene, in the 3’ UTR, 5’ UTR, introns, or exons. The number of mutation-carrying DE genes in the mutant lines ranged from 5 (0.43% of the total DE genes) to 423 (6.40%), with a median of 50.5 **(Supplementary Table S5)**. Furthermore, the ratio of down- *vs*. upregulated mutation-carrying DE genes ranged from 0.5 to 3, with mutant lines equally split between showing a majority of downregulated genes *vs* a majority of upregulated genes **(Figure 4B, Supplementary Table S5)**.

Except for lines RY-1 and RY-2 where ∼6% of DE genes carried a mutation, the remaining 14 lines all showed less than 2% of DE genes carrying a mutation **(Figure 4C)**. On the other hand, mutation-free DE genes in each mutant line ranged from 1171 (94.6%) to 6383 (99.6%) with a median of 3502 (**Figure 4C**). Therefore, with most DE genes being mutation-free, it strongly suggests that trans-effects drive the transcriptional changes of the majority of DE genes in these mutants and tend to result in more downregulation than upregulation, whereas cis-effects may only influence a small proportion of genes where trans-effects cannot be excluded from imposing their impacts. Lastly, cis mutations may not alter gene expression at all because we identified many genes carrying a mutation that did not show any differential expression. Out of the total number of genes carrying a mutation, the number of genes that were not differentially expressed ranged from 215 (51.6%) to 1021 (77.3%) across the lines, with a median of 303.

### Impact of EMS-induced mutations

Across all 16 mutant lines, the EMS-induced mutations were randomly distributed throughout the genome (chi-squared test, all p values > 0.05), i.e., the proportion of mutations in a specific type of genomic region does not exceed expectation based on random distribution of mutations. A total of 3047 (∼31%) mutations were found in the 5kb downstream region (5kb downstream of a gene), 3142 (∼32%) in the 5kb upstream region (5kb upstream of a gene), 1068 (∼11%) in exons, 1762 (∼18%) in intergenic regions, 538 (∼5%) in introns, 58 (∼0.6%) in splice sites, 144 (∼1.5%) in 3’ UTR, and 147 (∼1.5%) in 5’ UTR **(Figure 4D, Supplementary Table S6).**

Moreover, we analyzed the distribution of mutations located within differentially expressed (DE) genes, and mutations located in non-DE genes across all 16 mutant lines. The distribution of mutations in both categories of genes is highly similar. For the mutation-carrying DE genes, a total of 650 (∼34%) mutations were located in the 5kb downstream region, 698 (∼37%) in the 5kb upstream region, 311 (∼17%) in exons, 140 (∼7%) in introns, 16 (∼1%) in the splice site region, 43 (∼2%) in the 3’UTR, and 33 (∼2%) in the 5’UTR **(Figure 5A, Supplementary Table S7)**. On the other hand, for the mutation-carrying non-DE genes, 2397 (∼38%) in the 5kb downstream region, 2444 (∼39%) in the 5kb upstream region, 757 (∼12%) in exons, 398 (∼6%) in introns, 42 (∼1%) in splice site region, 101 (∼2%) in the 3’ UTR, and 114 (∼2%) in the 5’ UTR region **(Figure 5B, Supplementary Table S8)**.

**Figure 5.**
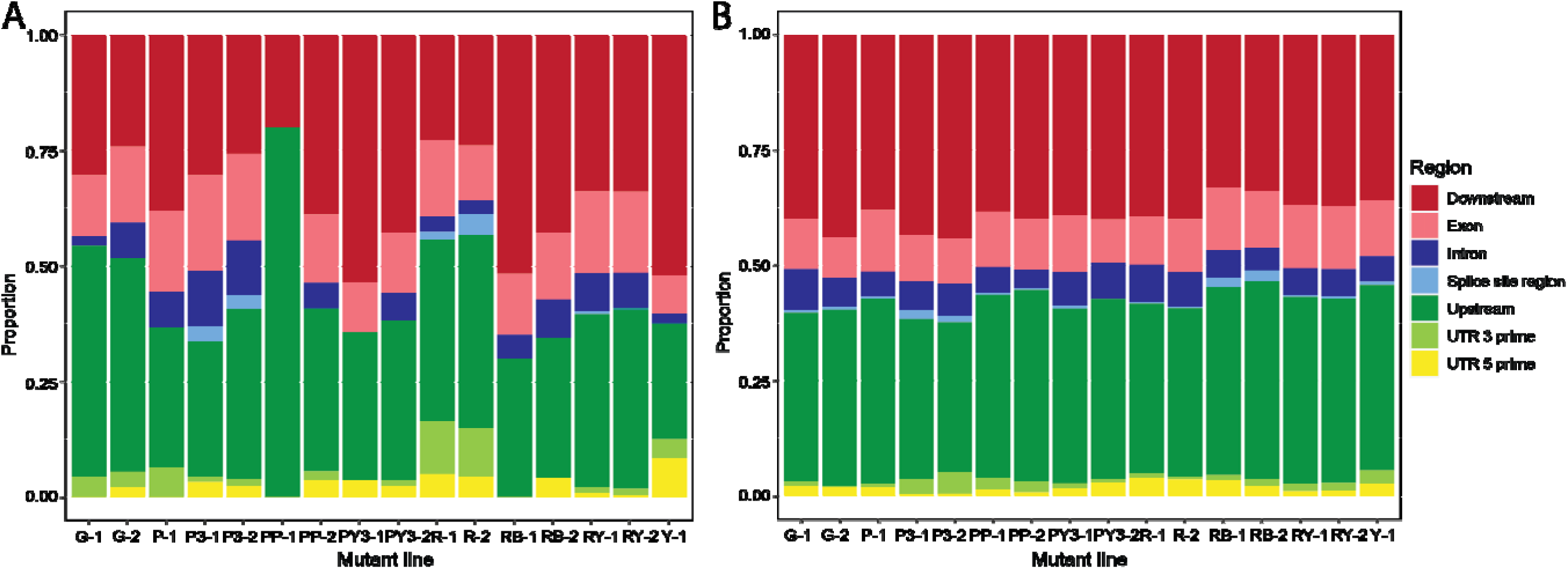
(A) Distribution of EMS-induced mutations in DE genes. (B) Distribution of EMS-induced mutations in non-DE genes.

The occurrence of mutation-carrying DE and non-DE genes presents an opportunity to see whether differential expression is associated with EMS-induced mutations in a specific genomic region. Using a multiple logistic regression model, we find that there was a significant association (p-value = 0.028) between differential expression and mutations in exons. Our results showed that for every mutation occurring in the exonic region of a gene, there is an increase of 0.73 in the log odds of that gene being differentially expressed, suggesting that exonic mutations are an important driver of altered gene expression.

Moreover, we categorized the mutations in DE and non-DE genes as high impact, moderate impact, low impact, or modifier variants. The mutations identified in DE genes consisted of 27 (1%) high impact variants (i.e., loss of a start codon, gain of a stop codon, splice acceptor variant), 184 (10%) moderate impact variants (i.e., missense variants), 113 (6%) low impact variants (i.e., splice region variant, synonymous variant, and variants producing a premature start codon in the 5’ UTR region), and 1553 (83%) modifier variants (i.e., variants 5kb upstream or downstream of a gene, variants within the 3’ UTR and 5’ UTR region, and variants within introns **(Supplementary Table S9)**.

Interestingly, for the high-impact variants, two genes carrying splice acceptor variants were both downregulated, while two other genes carrying start-lost variants were both upregulated. Out of 23 genes carrying stop-gained variants, only four were downregulated whereas the remaining 19 were upregulated **(Figure 6)**. From the genes carrying moderate impact (i.e., missense) variants, low impact variants and modifier variants; 129, 83 and 765 were upregulated, and 55, 36 and 793 were downregulated, respectively, suggesting moderate and low impact variants tend to be more associated with upregulated expression than downregulated expression **(Supplementary Table S10)**.

**Figure 6.**
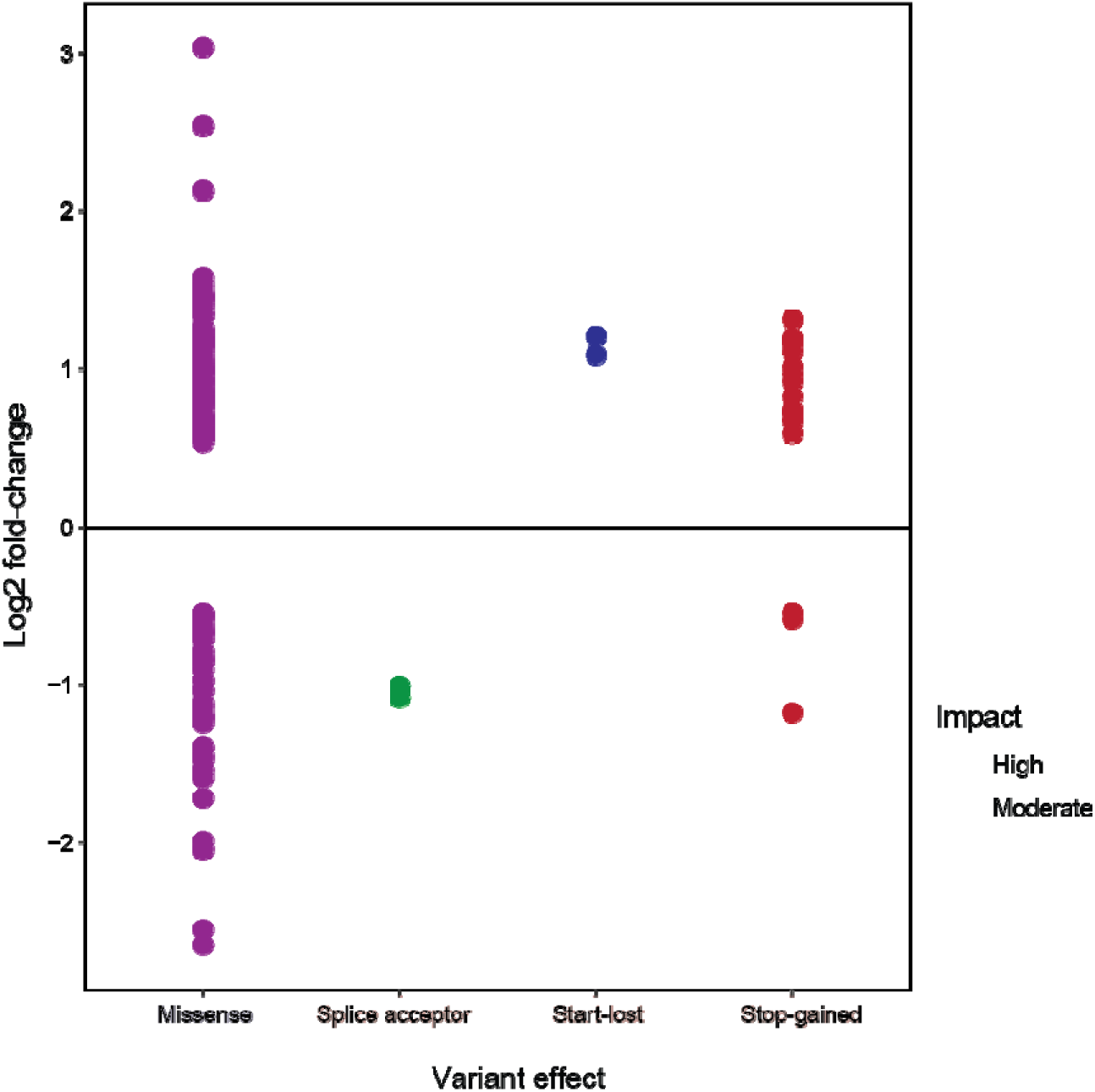
Log2 fold-change of DE genes with high-impact and moderate-impact variants identified across all 16 mutant lines.

On the other hand, mutations identified in non-DE genes have a similar distribution of the impact categories, with 32 (∼1%) high impact variants, 462 (∼7%) moderate impact variants, 303 (∼5%) low impact variants and 5441 (∼87%) modifier variants **(Supplementary Table S11 and S12)**. It is interesting to note that even some high-impact variants such as loss of a start codon and gain of a stop codon did not result in altered expression for the mutation carrying genes.

### Alternative splicing

Alternatively spliced (AS) genes were identified by comparing all 16 EMS-induced mutant lines to the wildtype. The number of alternatively spliced genes ranged from 212 to 627 across all mutant lines, with a median of 393 **(Supplementary Table S13)**. The number of mutation-carrying AS genes ranged from 2 (0.5% of the total AS genes) to 57 (9.6%) with a median of 12, while that of mutation-free AS genes ranged from 202 (∼90%) to 611 (∼99%) across mutant lines, with a median of 373 **(Figure 7, Supplementary Table S14 and 15)**. The dominance of mutation-free AS genes strongly suggests that the trans-effects caused by mutations are a major driver of alternative splicing.

**Figure 7.**
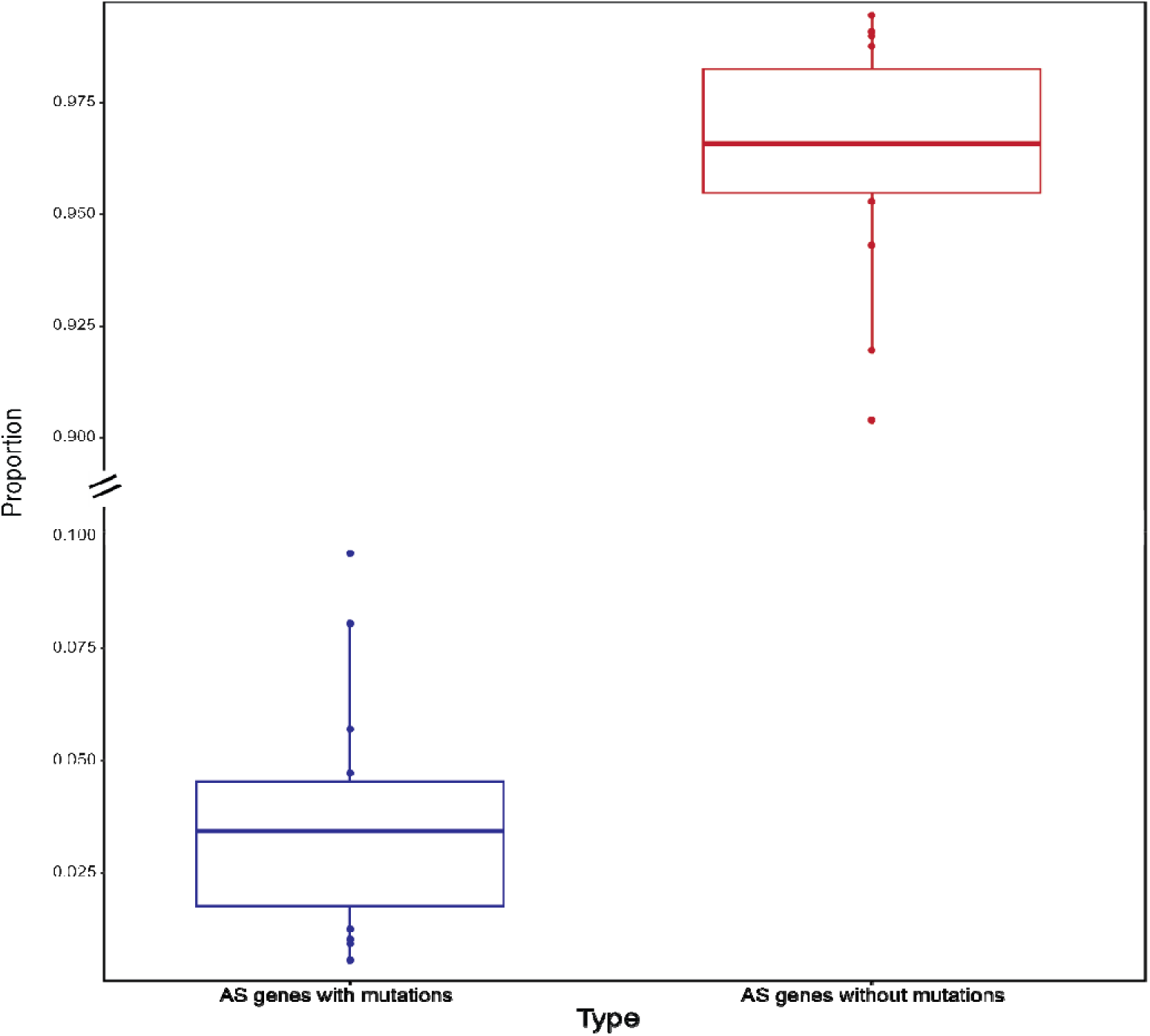
Proportion of alternatively spliced (AS) genes with EMS-induced mutations and AS genes without mutations.

Overall, the AS genes consist of 665 (11%) A3SS events, 665 (11%) A5SS events, 1051 (17%) MXE events, 2027 (32%) RI events, and 1880 (30%) SE events **(Supplementary Table S13)**. Specifically, mutation-carrying AS genes consisted of the following events: 20 (∼8%) A3SS, 22 (∼9%) A5SS, 48 (∼19%) MXE, 55 (∼22%) RI, and 104 (∼42%) SE **(Supplementary Table S14)**. On the other hand, mutation-free AS events were as the follows: 645 (∼11%) A3SS, 643 (∼11%) A5SS, 1003 (∼17%) MXE, 1972 (∼33%) RI and 1776 (∼29%) SE events **(Supplementary Table S15)**. No significant difference (chi-square test, p-value = 0.25) was observed in the distribution of AS events between mutation-carrying and mutation-free AS genes. Lastly, across all 16 EMS-induced mutant lines, a total of 44 genes were both AS and DE (median=2). Of these genes, 13 were downregulated and 31 were upregulated **(Supplementary Table S14)**.

### Impact and effect of genic EMS-induced mutations on alternative splicing

Mutations located in the alternatively spliced genes were further categorized according to their impact, with 6 (∼3%) being high impact, 14 (∼7%) low impact, 20 (∼10%) moderate impact, and 159 (∼80%) modifier impact variants across all 16 EMS-induced mutant lines **(Supplementary Table S16)**. The high impact variants consisted of 1 (∼0.5%) splice acceptor variant, 2 (∼1%) splice donor variant, and 3 (∼2%) stop-gained variants. Moderate impact variants consisted of 20 (∼10%) missense variants, while low impact variants consisted of 1 (∼0.5%) splice region variant and 13 (∼7%) synonymous variants. Modifier impact variants consisted of 1 (∼0.5%) 3’ UTR variant, 40 (∼20%) downstream variants, 22 (∼11%) intron variants, and 97 (∼49%) upstream variants across all mutant lines **(Supplementary Table S17)**. All alternatively spliced genes carried only one variant type, and only one gene from the P-1 mutant line carried 2 modifier mutations located in an intron and the 5kb upstream region. Lastly, results from a logistics regression showed that there was no significant association between AS event and variant type. However, downstream variants (p-value = 0.080), missense variants (p-value 0.053) and upstream variants (p-value = 0.078) were marginally significant, while all other variant types have p-values > 0.99.

## Discussion

How mutations alter genome-wide gene expression and alternative splicing has been understudied. Only a limited number of studies have attempted to address this issue using mutation-accumulation experiments in a small number of species including *C. elegans*, *D. melanogaster*, *S. cerevisiae,* and zebrafish (Denver *et al*. 2005, Huang *et al*. 2016, Kim *et al*. 2007, Konrad *et al*. 2018, Landry *et al*. 2007, Rifkin *et al*. 2005, White *et al*. 2022, Zalts and Yanai 2017). In this study, we examined the alterations in genome-wide gene expression and alternative splicing patterns in EMS-induced mutants of the microcrustacean *Daphnia* to bridge this knowledge gap. Furthermore, we dissected the contribution of cis and trans mutations to differential gene expression (DE) and alternative splicing (AS).

### Global gene expression changes due to EMS-induced mutations

In our mutant lines the EMS-induced base substitution rate, per gene mutations rate, and non-synonymous mutation rate were estimated to be (medians) 1.84×10^-6^ per site per generation, 3.1×10^-3^ per gene per generation and 1.14×10^-3^ per gene per generation respectively **(Figure 3A, B)**. These estimates are derived from a rigorous mutation filtering pipeline that has been calibrated with PCR verification of mutations in a previous study (Snyman *et al*. 2021), with a false discovery rate <0.05. These rate estimates are also consistent with our previous work (Snyman *et al*. 2021), showing elevated mutation rates hundreds of times higher than the spontaneous mutation rates in *Daphnia* (Bull *et al*. 2019, Flynn *et al*. 2017, Keith *et al*. 2016). The collection of identified mutations in these mutant lines show the notable signature of EMS-induced mutations, i.e., significant enrichment of G:C to A:T transitions **(Figure 3C)** and a random distribution across the genome **(Figure 4A**, Greene *et al*. 2003).

Considering that EMS may cause acute gene expression changes in the exposed females and the lingering maternal effects may alter gene expression in immediate offspring, we measured gene expression in the clonal mutant lines several generations after the exposure. This method ensures that the observed gene expression changes are most likely caused by the EMS-induced mutations, rather than by maternal effects.

Our differential expression analyses on the mutant lines reveal that downregulated genes are more abundant than upregulated genes in nearly all mutant lines, with the ratio of down- *vs.* upregulated genes ranging between 1.05 and 3.63 **(Figure 4B, Supplementary Table S5)**. It is not clear whether mutations induced by other mutagens would cause a similar pattern as gene expression changes in mutagen-induced mutant lines remains understudied. However, as we discuss below the dominance of trans effects in gene expression changes, the higher abundance of downregulation is likely caused by trans-acting mutations mainly acting as silencers to reduce transcription (Johnson *et al*. 2015).

Examining whether differentially expressed (DE) genes carry EMS-induced mutations across our mutant lines unveiled a significant contribution of trans mutations to expression changes. DE genes carrying EMS-induced mutations only accounted for a small portion of the total (ranging from 0.43% to 6.40% across our mutant lines), while mutation-free DE genes showed dominance (ranging from 94.6% to 99.6% across mutant lines). This pattern strongly suggests that the pleiotropic effects of trans mutations are a major contributor to the variance in gene expression between mutant and wildtype lines **(Supplementary Table S5),** while cis mutations only play a relatively small role. The pleiotropic effects of trans mutations were previously reported in *D. melanogaster* and *C. elegans* mutation accumulation lines, where a relatively small number of trans-acting mutations with multiple downstream effects induced the majority of the transcriptional differences (Denver *et al*. 2005, Huang *et al*. 2016). The dominant pleiotropic effects of trans mutations are probably due to trans factors representing a larger mutational target and are more likely to arise than cis mutations (Chen *et al*. 2015, Coolon *et al*. 2014, Gruber *et al*. 2012, Metzger *et al*. 2016, Rhoné *et al*. 2017, Wittkopp *et al*. 2004).

The presence of cis mutations in DE genes indicates that cis mutations may directly cause gene expression changes. However, it cannot be ruled out that trans-effects also contribute to the variance in gene expression of these genes. It is noteworthy that the presence of cis mutations does not necessarily mean altered expression because 51.6% - 77.3% of the observed genes with cis mutations do not show differential expression. However, our logistic regression analysis provides a clear signal that exonic mutations are strongly associated with altered gene expression. Interestingly, it was observed in EMS mutagenized *S. cerevisiae* mutants that trans regulatory mutations were also mostly located within the coding sequences (Duveau *et al*. 2021).

It is possible that some cis mutations do not directly impact gene expression. However, the unchanged expression of genes with cis mutations may also be a result of complex interactions between cis and trans effects, for which we do not have the capacity to further test in this study. Furthermore, because the mutated sites are heterozygous for the mutation and wildtype SNP allele, the invariance of total gene expression may have concealed the signature of allele-specific expression changes caused by the EMS-induced mutations. It would be of great interest for future studies to address whether and how mutations cause the changes in allele-specific transcript abundance.

### Impact of different classes of mutations on gene expression

Our analyses yielded insights into the distribution and impact of different classes of mutations on gene expression. The distribution of mutations in DE and non-DE genes were highly similar with the majority found in the 5kb upstream and downstream regions **(Figure 5A and B, Supplementary Table S7 and S8)**. Mutations identified in DE genes were mostly modifier impact variants (i.e., variants 5kb upstream or downstream of a gene, variants within the 3’ UTR and 5’ UTR region, and variants within introns) followed by moderate impact (i.e., missense variants), low impact (i.e., splice region variant, synonymous variant, and variants producing a premature start codon in the 5’ UTR region) and high impact (i.e., loss of a start codon, gain of a stop codon, splice acceptor variant) variants **(Supplementary Table S9)**. Interestingly, our results also showed that moderate-impact (missense) and low-impact variants tend to be associated with an upregulation of gene expression (129 upregulated genes vs 55 downregulated genes for moderate impact, 83 upregulated vs 36 downregulated genes for low impact, **Figure 6)**. Missense variants are known to alter the amino acid sequence, and have been shown to impact the interacting DNA-transcription factors, resulting in changed expression of other genes (Ding *et al*. 2015). Low-impact variants including synonymous variants have also been shown to influence gene expression by disrupting transcription and splicing, as well as mRNA stability (Pagani *et al*. 2005, Presnyak *et al*. 2015, Stergachis *et al*. 2013, Wang *et al*. 2021).

High impact variants consisted of two downregulated splice acceptor variants, two upregulated start-lost variants, and the upregulation of most stop-gained variants **(Figure 6)**. The downregulation of some stop-gained (nonsense) variants could indicate nonsense-mediated RNA decay (NMD) is at play. NMD is a surveillance pathway that helps maintain RNA quality and cellular homeostasis by detecting and eliminating transcripts containing premature stop codons (Nickless *et al*. 2017). The upregulation of most stop-gained variants could indicate that these genes escape NMD, that a stop-codon read-through occurs, or that transcriptional adaptation is triggered causing an upregulation of the affected as well as related genes (El-Brolosy *et al*. 2019).

On the other hand, mutation-carrying non-DE genes showed a similar distribution to DE genes. Interestingly, some high-impact mutations in these genes did not result in an altered expression **(Supplementary Table S11 and S12)**. It is plausible that some of these mutations are false positives that slipped through our mutation identification pipeline, although the rate of false positives should be < 5%. Another possibility is that our RNA-seq experimental design did not provide enough experimental power to detect the expression changes of these genes. It should also be noted that only one allele of the affected genes was mutated, which means that a functional wildtype allele still exists for the affected genes. Thus, it is possible that while the mutated allele may have had reduced transcription, the wildtype allele likely experiences enhanced transcription, resulting in unchanged expression of the gene in question.

### Alternative splicing

Changes in pre-mRNA splicing are another important source of phenotypic variation. Exon-intron boundaries consist of highly conserved splice site sequences that must be correctly identified by the spliceosome to perform a splicing reaction. Additionally, splicing factors interact with intronic and exonic sequences to determine how frequently an exon is included in the final transcript (Wang and Burge 2008). If the splicing process is not correctly regulated, premature stop codons and transcripts with an altered amino acid sequence can result. Alternative splicing (AS) can result from both cis- and trans-regulatory mutations. Cis mutations can interact with exonic or intronic splicing enhancers or silencers, whereas transmutations affect splicing factors that can impact genes throughout the genome (Juan *et al*. 2014, Kornblihtt *et al*. 2013). To date, the impact of cis versus trans mutations on alternative splicing remains understudied. Most studies have suggested that cis-regulatory (Ast 2004, Keren *et al*. 2010, Thatcher *et al*. 2014) mutations may be the main contributor to alternative splicing. However, a recent study raised the opposite view, suggesting that the pleiotropic effects of trans-mutations may be the main culprit (Smith *et al*. 2018).

Our results are consistent with the latter view. Our analyses show that the number of mutation-carrying AS genes ranged from 0.5% to 9.6% across mutant lines **(Figure 7, Supplementary Table S13)**. Most AS genes are free of EMS-induced mutations, (ranging from 90% to 99% of total AS genes), strongly suggesting that trans-effects are a major driver of alternative splicing **(Figure 7, Supplementary Table S14 and S15**). Furthermore, our results showed that there was little overlap between differentially expressed and alternatively spliced genes, a pattern also seen in *Drosophila* (Jakšić and Schlötterer 2016), aphids (Grantham and Brisson 2018), and salmonids (Jacobs and Elmer 2021).

Our logistics regression analysis showed that there was no significant association between AS events and any variant type. However, downstream variants (p-value = 0.080), missense variants (p-value 0.053) and upstream variants (p-value = 0.078) were marginally significant, while all other variant types have p-values > 0.99. Mutations from these regions warrant further investigation in future studies.

### Implications for understanding the biological impacts of anthropogenic events

As we are preparing this manuscript, on February 3^rd^ 2023, a train derailment accident in East Palestine, Ohio, USA, resulted in the spill of > 100,000 gallons of vinyl chloride, a carcinogenic mutagen. Accidents like this and many unreported, smaller-scale anthropogenic events occur at a sharply increased rate since the Industrial Revolution. The epigenetic effects of some of these environmental chemicals have been investigated (reviewed in Baccarelli and Bollati 2009, Goyal *et al*. 2022, Hou *et al*. 2012). However, much work still needs to be done to determine the impact of chemically induced mutations on gene expression and alternative splicing. Based on the results from our study, mutations exerting trans effects can have a genome-wide impact by altering the expression and splicing of thousands of genes (up to 99% of the total differentially expressed and alternatively spliced genes respectively in *Daphnia*), while high impact and moderate impact mutations can lead to phenotypic alterations by changing the amino acid sequence. Understanding the consequences of these anthropogenic stressors are thus vital to protecting and preserving the health of our ecosystems and ourselves.

## Conclusions

Our results show that trans effects are the major contributor to the variance in gene expression and alternative splicing between the wildtype and EMS-induced mutant lines in *Daphnia*, while cis mutations do not always alter total gene expression and only affected a small portion of genes. Furthermore, our results showed a significant association between DE genes and exonic mutations, indicating that exonic mutations are an important driver of altered gene expression.

## Acknowledgments

We thank A. Gamadia and M.T. Smith for their help with tissue collection and Trung Huynh for his help with writing the Python scripts. Additionally, we would like to thank all Xu lab members for their helpful discussions. This work is supported by NIH grant R35GM133730 and NSF CAREER grant MCB-2042490 to SX.

## Author Contributions

SX designed the study, and MS and SX wrote the manuscript. MS performed the tissue collection, molecular work, and data analysis.

## References

Abzhanov A, Kuo WP, Hartmann C, Grant BR, Grant PR, Tabin CJ (2006) The calmodulin pathway and evolution of elongated beak morphology in Darwin’s finches. Nature, 442(7102), 563–567.

Albert FW, Bloom JS, Siegel J, Day L, Kruglyak L (2018) Genetics of trans-regulatory variation in gene expression. ELife, 7, e35471.

Anna A, Monika G (2018) Splicing mutations in human genetic disorders: Examples, detection, and confirmation. Journal of Applied Genetics, 59(3), 253–268.

Aquadro CF, Tachida H, Langley CH, Harada K, Mukai T (1990) Increased variation in ADH enzyme activity in Drosophila mutation-accumulation experiment is not due to transposable elements at the Adh structural gene. Genetics, 126(4), 915–919.

Ast G (2004) How did alternative splicing evolve? Nature Reviews. Genetics, 5(10), 773–782.

Andrews S (2010) FastQC: A Quality Control Tool for High Throughput Sequence Data [Online]. Available online at: http://www.bioinformatics.babraham.ac.uk/projects/fastqc/

Baccarelli A, Bollati V (2009) Epigenetics and environmental chemicals. Current Opinion in Pediatrics, 21(2), 243–251.

Bolger AM, Lohse M, Usadel B (2014) Trimmomatic: A flexible trimmer for Illumina sequence data. Bioinformatics (Oxford, England), 30(15), 2114–2120.

Bull JK, Flynn JM, Chain FJJ, Cristescu ME (2019) Fitness and Genomic Consequences of Chronic Exposure to Low Levels of Copper and Nickel in *Daphnia pulex* Mutation Accumulation Lines. G3: Genes, Genomes, Genetics, 9(1), 61–71.

Chen J, Nolte V, Schlötterer C (2015) Temperature stress mediates decanalization and dominance of gene expression in Drosophila melanogaster. PLoS Genetics, 11(2), e1004883.

Chu XL, Zhang QG (2021) Consequences of mutation accumulation for growth performance are more likely to be resource-dependent at higher temperatures. BMC Ecology and Evolution, 21(1), 109.

Cingolani P, Platts A, Wang LL, Coon M, Nguyen T, Wang L, Land SJ, Lu X, Ruden DM (2012) A program for annotating and predicting the effects of single nucleotide polymorphisms, SnpEff. Fly, 6(2), 80–92.

Colbourne JK, Pfrender ME, Gilbert D, Thomas WK, Tucker A, Oakley TH, Tokishita S, et al. (2011) The ecoresponsive genome of *Daphnia pulex*. Science (New York, N.Y.), 331(6017), 555–561.

Coolon JD, McManus CJ, Stevenson KR, Graveley BR, Wittkopp PJ (2014) Tempo and mode of regulatory evolution in *Drosophila*. Genome Research, 24(5), 797–808.

Coolon JD, Stevenson KR, McManus CJ, Yang B, Graveley BR, Wittkopp PJ (2015) Molecular Mechanisms and Evolutionary Processes Contributing to Accelerated Divergence of Gene Expression on the *Drosophila* X Chromosome. Molecular Biology and Evolution, 32(10), 2605–2615.

Denkena J, Johannes F, Colomé-Tatché M (2021) Region-level epimutation rates in Arabidopsis thaliana. Heredity, 127(2), Article 2.

Denver DR, Dolan PC, Wilhelm LJ, Sung W, Lucas-Lledó JI, Howe DK, Lewis SC, Okamoto K, Thomas WK, Lynch M, Baer CF (2009) A genome-wide view of *Caenorhabditis elegans* base-substitution mutation processes. Proceedings of the National Academy of Sciences, 106(38), 16310–16314.

Denver DR, Morris K, Streelman JT, Kim SK, Lynch M, Thomas WK (2005) The transcriptional consequences of mutation and natural selection in *Caenorhabditis elegans*. Nature Genetics, 37(5), 544–548.

Díaz-González J, Domínguez A (2020) Different structural variants of roo retrotransposon are active in *Drosophila melanogaster*. Gene, 741, 144546.

Ding J, McConechy MK, Horlings HM, Ha G, Chun Chan F, Funnell T, Mullaly SC, Reimand J, Bashashati A, Bader GD, Huntsman D, Aparicio S, Condon A, Shah SP (2015) Systematic analysis of somatic mutations impacting gene expression in 12 tumour types. Nature Communications, 6, 8554.

Dobin A, Davis CA, Schlesinger F, Drenkow J, Zaleski C, Jha S, Batut P, Chaisson M, Gingeras TR (2013) STAR: Ultrafast universal RNA-seq aligner. Bioinformatics, 29(1), 15–21.

Doyle JJ, Doyle JL (1987) A rapid DNA isolation procedure for small quantities of fresh leaf tissue (RESEARCH). PHYTOCHEMICAL BULLETIN.

Duveau F, Vande Zande P, Metzger BP, Diaz CJ, Walker EA, Tryban S, Siddiq MA, Yang B, Wittkopp PJ (2021) Mutational sources of trans-regulatory variation affecting gene expression in *Saccharomyces cerevisiae*. ELife, 10, e67806.

Edison AS, Hall RD, Junot C, Karp PD, Kurland IJ, Mistrik R, Reed LK, Saito K, Salek RM, Steinbeck C, Sumner LW, Viant, MR (2016) The Time Is Right to Focus on Model Organism Metabolomes. Metabolites, 6(1), 8.

El-Brolosy MA, Kontarakis Z, Rossi A, Kuenne C, Günther S, Fukuda N et al. (2019) Genetic compensation triggered by mutant mRNA degradation. Nature, 568(7751), 193–197.

Emerson JJ, Hsieh LC, Sung HM, Wang TY, Huang CJ, Lu HHS, Lu MYJ, Wu SH, Li WH (2010) Natural selection on cis and trans regulation in yeasts. Genome Research, 20(6), 826–836.

Flibotte S, Edgley ML, Chaudhry I, Taylor J, Neil SE, Rogula A, Zapf R, Hirst M, Butterfield Y, Jones SJ, Marra MA, Barstead RJ, Moerman DG (2010) Whole-Genome Profiling of Mutagenesis in *Caenorhabditis elegans*. Genetics, 185(2), 431–441.

Flynn JM, Caldas I, Cristescu ME, Clark AG (2017) Selection Constrains High Rates of Tandem Repetitive DNA Mutation in *Daphnia pulex*. Genetics, 207(2), 697–710.

Flynn JM, Chain FJJ, Schoen DJ, Cristescu ME (2017) Spontaneous Mutation Accumulation in *Daphnia pulex* in Selection-Free vs. Competitive Environments. Molecular Biology and Evolution, 34(1), 160–173.

Frisch D, Morton PK, Chowdhury PR, Culver BW, Colbourne JK, Weider LJ, Jeyasingh PD (2014) A millennial-scale chronicle of evolutionary responses to cultural eutrophication in *Daphnia*. Ecology Letters, 17(3), 360–368.

Ghalambor CK, Hoke KL, Ruell EW, Fischer EK, Reznick DN, Hughes KA (2015) Non-adaptive plasticity potentiates rapid adaptive evolution of gene expression in nature. Nature, 525(7569), 372–375.

Gorr TA, Rider CV, Wang HY, Olmstead AW, LeBlanc GA (2006) A candidate juvenoid hormone receptor cis-element in the *Daphnia magna* hb2 hemoglobin gene promoter. Molecular and Cellular Endocrinology, 247(1–2), 91–102.

Goyal K, Goel H, Baranwal P, Dixit A, Khan F, Jha NK, Kesari KK, Pandey P, Pandey A, Benjamin M, Maurya A, Yadav V, Sinh RS, Tanwar P, Upadhyay TK, Mittan S (2022) Unravelling the molecular mechanism of mutagenic factors impacting human health. Environmental Science and Pollution Research, 29(41), 61993–62013.

Grantham ME, Brisson JA (2018) Extensive Differential Splicing Underlies Phenotypically Plastic Aphid Morphs. Molecular Biology and Evolution, 35(8), 1934–1946.

Graur D, Li WH (2000) Fundamentals of Molecular Evolution (2nd Edition). Sinauer Associates. Sunderland MA

Greene EA, Codomo CA, Taylor NE, Henikoff JG, Till BJ, Reynolds SH, Enns LC, Burtner C, Johnson JE, Odden AR, Comai L, Henikoff S (2003) Spectrum of chemically induced mutations from a large-scale reverse-genetic screen in *Arabidopsis*. Genetics, 164(2), 731–740.

Gruber JD, Vogel K, Kalay G, Wittkopp PJ (2012) Contrasting properties of gene-specific regulatory, coding, and copy number mutations in *Saccharomyces cerevisiae*: Frequency, effects, and dominance. PLoS Genetics, 8(2), e1002497.

Harada K (1995) A quantitative analysis of modifier mutations which occur in mutation accumulation lines in Drosophila melanogaster. Heredity, 75(6), Article 6.

Ho EKH, Bellis ES, Calkins J, Adrion JR, Iv LCL, Schaack S (2021) Engines of change: Transposable element mutation rates are high and variable within Daphnia magna. PLOS Genetics, 17(11), e1009827.

Hornung G, Oren M, Barkai N (2012) Nucleosome organization affects the sensitivity of gene expression to promoter mutations. Molecular Cell, 46(3), 362–368.

Hou L, Zhang X, Wang D, Baccarelli A (2012) Environmental chemical exposures and human epigenetics. International Journal of Epidemiology, 41(1), 79–105.

Huang W, Lyman RF, Lyman RA, Carbone MA, Harbison ST, Magwire MM, Mackay TF (2016) Spontaneous mutations and the origin and maintenance of quantitative genetic variation. ELife, 5, e14625.

Jackson CE, Xu S, Ye Z, Pfrender ME, Lynch M, Colbourne JK, Shaw JR (2021) Chromosomal rearrangements preserve adaptive divergence in ecological speciation (p. 2021.08.20.457158).

Jacobs A, Elmer KR (2021) Alternative splicing and gene expression play contrasting roles in the parallel phenotypic evolution of a salmonid fish. Molecular Ecology, 30(20), 4955– 4969.

Jakšić AM, Schlötterer C (2016) The Interplay of Temperature and Genotype on Patterns of Alternative Splicing in *Drosophila melanogaster*. Genetics, 204(1), 315–325.

Jiang C, Mithani A, Belfield EJ, Mott R, Hurst LD, Harberd NP (2014) Environmentally responsive genome-wide accumulation of de novo *Arabidopsis thaliana* mutations and epimutations. Genome Research, 24(11), 1821–1829.

Johnson WC, Ordway AJ, Watada M, Pruitt JN, Williams TM, Rebeiz M (2015) Genetic Changes to a Transcriptional Silencer Element Confers Phenotypic Diversity within and between *Drosophila* Species. PLoS Genetics, 11(6), e1005279.

Josephs EB (2021) Gene expression links genotype and phenotype during rapid adaptation. Molecular Ecology, 30(1), 30–32.

Juan WC, Roca X, Ong ST (2014) Identification of cis-Acting Elements and Splicing Factors Involved in the Regulation of BIM Pre-mRNA Splicing. PLOS ONE, 9(4), e95210.

Keightley PD, Trivedi U, Thomson M, Oliver F, Kumar S, Blaxter ML (2009) Analysis of the genome sequences of three *Drosophila melanogaster* spontaneous mutation accumulation lines. Genome Research, 19(7), 1195–1201.

Keith N, Tucker AE, Jackson CE, Sung W, Lucas Lledó JI, Schrider DR, Schaack S, Dudycha JL, Ackerman M, Younge AJ, Shaw J R, Lynch M (2016) Generation of loss- or gain of function mutants and weak nonlethal alleles. Genome Research, 26(1), 60–69.

Keren H, Lev-Maor G, Ast G (2010) Alternative splicing and evolution: Diversification, exon definition and function. Nature Reviews. Genetics, 11(5), 345–355.

Kilham SS, Kreeger DA, Lynn SG, Goulden CE, Herrera L (1998) COMBO: A defined freshwater culture medium for algae and zooplankton. Hydrobiologia, 377(1), 147–159.

Kim N, Abdulovic AL, Gealy R, Lippert MJ, Jinks-Robertson S (2007) Transcription-associated mutagenesis in yeast is directly proportional to the level of gene expression and influenced by the direction of DNA replication. DNA Repair, 6(9), 1285–1296.

Konrad A, Flibotte S, Taylor J, Waterston RH, Moerman DG, Bergthorsson U, Katju V (2018) Mutational and transcriptional landscape of spontaneous gene duplications and deletions in *Caenorhabditis elegans*. Proceedings of the National Academy of Sciences of the United States of America, 115(28), 7386–7391.

Kornblihtt AR, Schor IE, Alló M, Dujardin G, Petrillo E, Muñoz MJ (2013) Alternative splicing: A pivotal step between eukaryotic transcription and translation. Nature Reviews Molecular Cell Biology, 14(3), Article 3.

Krasovec M (2021) The spontaneous mutation rate of *Drosophila pseudoobscura*. G3 Genes|Genomes|Genetics, 11(7), jkab151.

Kwasnieski JC, Mogno I, Myers CA, Corbo JC, Cohen BA (2012) Complex effects of nucleotide variants in a mammalian cis-regulatory element. Proceedings of the National Academy of Sciences of the United States of America, 109(47), 19498–19503.

Landrigan PJ, Fuller R, Acosta NJR, Adeyi O, Arnold R, Basu N, Baldé AB, Bertollini R, Bose-O’Reilly S, Boufford J. I, Breysse PN et al. (2018) The Lancet Commission on pollution and health. The Lancet, 391(10119), 462–512.

Landry CR, Lemos B, Rifkin SA, Dickinson WJ, Hartl DL (2007) Genetic properties influencing the evolvability of gene expression. Science (New York, N.Y.), 317(5834), 118–121.

Latta LC, Peacock M, Civitello DJ, Dudycha JL, Meik JM, Schaack S, Baer AECF, Kalisz ES (2015) The Phenotypic Effects of Spontaneous Mutations in Different Environments. The American Naturalist, 185(2), 243–252.

Lee SY, Cheong JI, Kim TS (2003) Production of doubled haploids through anther culture of M1 rice plants derived from mutagenized fertilized egg cells. Plant Cell Reports, 22(3), 218– 223.

Lewis JA, Broman AT, Will J, Gasch AP (2014) Genetic architecture of ethanol-responsive transcriptome variation in *Saccharomyces cerevisiae* strains. Genetics, 198(1), 369–382.

Li H (2011) A statistical framework for SNP calling, mutation discovery, association mapping and population genetical parameter estimation from sequencing data. Bioinformatics, 27(21), 2987–2993.

Li H, Durbin R (2009) Fast and accurate short read alignment with Burrows-Wheeler transform. Bioinformatics (Oxford, England), 25(14), 1754–1760.

Li H, Handsaker B, Wysoker A, Fennell T, Ruan J, Homer N, Marth G, Abecasis G, Durbin R, 1000 Genome Project Data Processing Subgroup (2009) The Sequence Alignment/Map format and SAMtools. Bioinformatics (Oxford, England), 25(16), 2078–2079.

Liao Y, Smyth GK, Shi W (2014) featureCounts: An efficient general purpose program for assigning sequence reads to genomic features. Bioinformatics (Oxford, England), 30(7), 923–930.

Love MI, Huber W, Anders S (2014) Moderated estimation of fold change and dispersion for RNA-seq data with DESeq2. Genome Biology, 15(12), 550.

Luu DHK, Vo HP, Xu S (2020) An efficient method for hatching diapausing embryos of *Daphnia pulex* species complex (Crustacea, Anomopoda). Journal of Experimental Zoology Part A: Ecological and Integrative Physiology, 333(2), 111–117.

Mack KL, Ballinger MA, Phifer-Rixey M, Nachman MW (2018) Gene regulation underlies environmental adaptation in house mice. Genome Research, 28(11), 1636–1645.

Macknight R, Duroux M, Laurie R, Dijkwel P, Simpson G, Dean C (2002) Functional significance of the alternative transcript processing of the Arabidopsis floral promoter FCA. The Plant Cell, 14(4), 877–888.

Manceau M, Domingues VS, Mallarino R, Hoekstra HE (2011) The developmental role of Agouti in color pattern evolution. Science (New York, N.Y.), 331(6020), 1062–1065.

Maricque BB, Dougherty JD, Cohen BA (2017) A genome-integrated massively parallel reporter assay reveals DNA sequence determinants of cis-regulatory activity in neural cells. Nucleic Acids Research, 45(4), e16.

Melnikov A, Murugan A, Zhang X, Tesileanu T, Wang L, Rogov P, Feizi S, Gnirke A et al. (2012) Systematic dissection and optimization of inducible enhancers in human cells using a massively parallel reporter assay. Nature Biotechnology, 30(3), 271–277.

Metzger BPH, Duveau F, Yuan DC, Tryban S, Yang B, Wittkopp PJ (2016) Contrasting Frequencies and Effects of cis- and trans-Regulatory Mutations Affecting Gene Expression. Molecular Biology and Evolution, 33(5), 1131–1146.

Metzger BPH, Yuan DC, Gruber JD, Duveau F, Wittkopp PJ (2015) Selection on noise constrains variation in a eukaryotic promoter. Nature, 521(7552), 344–347.

Nickless A, Bailis JM, You Z (2017) Control of gene expression through the nonsense-mediated RNA decay pathway. Cell & Bioscience, 7(1), 26.

Omilian AR, Cristescu MEA, Dudycha JL, Lynch M (2006) Ameiotic recombination in asexual lineages of *Daphnia*. Proceedings of the National Academy of Sciences of the United States of America, 103(49), 18638–18643.

Ossowski S, Schneeberger K, Lucas-Lledó JI, Warthmann N, Clark RM, Shaw RG, Weigel D, Lynch M (2010) The Rate and Molecular Spectrum of Spontaneous Mutations in *Arabidopsis thaliana*. Science (New York, N.Y.), 327(5961).

Pagani F, Raponi M, Baralle FE (2005) Synonymous mutations in CFTR exon 12 affect splicing and are not neutral in evolution. Proceedings of the National Academy of Sciences, 102(18), 6368–6372.

Pastink A, Heemskerk E, Nivard MJ, van Vliet CJ, Vogel EW (1991) Mutational specificity of ethyl methanesulfonate in excision-repair-proficient and -deficient strains of *Drosophila melanogaster*. Molecular & General Genetics: MGG, 229(2), 213–218.

Patwardhan RP, Lee C, Litvin O, Young DL, Pe’er D, Shendure J (2009) High-resolution analysis of DNA regulatory elements by synthetic saturation mutagenesis. Nature Biotechnology, 27(12), Article 12.

Pohl M, Bortfeldt RH, Grützmann K, Schuster S (2013) Alternative splicing of mutually exclusive exons—A review. Biosystems, 114(1), 31–38.

Presnyak V, Alhusaini N, Chen YH, Martin S, Morris N, Kline N, Olson S, Weinberg D, Baker KE, Graveley BR, Coller J (2015) Codon Optimality Is a Major Determinant of mRNA Stability. Cell, 160(6), 1111–1124.

Rebeiz M, Pool JE, Kassner VA, Aquadro CF, Carroll SB (2009) Stepwise modification of a modular enhancer underlies adaptation in a *Drosophila* population. Science (New York, N.Y.), 326(5960), 1663–1667.

Rhoné B, Mariac C, Couderc M, Berthouly-Salazar C, Ousseini IS, Vigouroux Y (2017) No Excess of Cis-Regulatory Variation Associated with Intraspecific Selection in Wild Pearl Millet (Cenchrus americanus). Genome Biology and Evolution, 9(2), 388–397.

Rifkin SA, Houle D, Kim J, White KP (2005) A mutation accumulation assay reveals a broad capacity for rapid evolution of gene expression. Nature, 438(7065), Article 7065.

Schaefke B, Emerson JJ, Wang TY, Lu MYJ, Hsieh LC, Li WH (2013) Inheritance of gene expression level and selective constraints on trans- and cis-regulatory changes in yeast. Molecular Biology and Evolution, 30(9), 2121–2133.

Scheffer H, Coate JE, Ho EKH, Schaack S (2022) Thermal stress and mutation accumulation increase heat shock protein expression in *Daphnia*. Evolutionary Ecology, 36(5), 829– 844.

Sharon E, Kalma Y, Sharp A, Raveh-Sadka T, Levo M, Zeevi D, Keren L, Yakhini Z, Weinberger A, Segal E (2012) Inferring gene regulatory logic from high-throughput measurements of thousands of systematically designed promoters. Nature Biotechnology, 30(6), 521–530.

Shen S, Park JW, Lu Z, Lin L, Henry MD, Wu YN, Zhou Q, Xing Y (2014) rMATS: Robust and flexible detection of differential alternative splicing from replicate RNA-Seq data. Proceedings of the National Academy of Sciences of the United States of America, 111(51), E5593–5601.

Smith CCR, Tittes S, Mendieta JP, Collier-zans E, Rowe HC, Rieseberg LH, Kane NC (2018) Genetics of alternative splicing evolution during sunflower domestication. Proceedings of the National Academy of Sciences, 115(26), 6768–6773.

Snyman M, Huynh TV, Smith MT, Xu S (2021) The genome-wide rate and spectrum of EMS-induced heritable mutations in the microcrustacean *Daphnia*: On the prospect of forward genetics. Heredity, 127(6), 535–545.

Stergachis AB, Haugen E, Shafer A, Fu W, Vernot B, Reynolds A, Raubitschek A, Ziegler S, LeProust EM, Akey JM, Stamatoyannopoulos JA (2013) Exonic Transcription Factor Binding Directs Codon Choice and Affects Protein Evolution. Science, 342(6164), 1367– 1372.

Suresh S, Crease TJ, Cristescu ME, Chain FJJ (2020) Alternative splicing is highly variable among *Daphnia pulex* lineages in response to acute copper exposure. BMC Genomics, 21(1), 433.

Tatarazako N, Oda S, Watanabe H, Morita M, Iguchi T (2003) Juvenile hormone agonists affect the occurrence of male *Daphnia*. Chemosphere, 53(8), 827–833.

Terai Y, Morikawa N, Kawakami K, Okada N (2003) The complexity of alternative splicing of hagoromo mRNAs is increased in an explosively speciated lineage in East African cichlids. Proceedings of the National Academy of Sciences of the United States of America, 100(22), 12798–12803.

Thatcher SR, Zhou W, Leonard A, Wang BB, Beatty M, Zastrow-Hayes G, Zhao X, Baumgarten A, Li B (2014) Genome-wide analysis of alternative splicing in Zea mays: Landscape and genetic regulation. The Plant Cell, 26(9), 3472–3487.

Vande Zande P, Hill MS, Wittkopp PJ (2022) Pleiotropic effects of trans-regulatory mutations on fitness and gene expression. Science, 377(6601), 105–109.

Vernia S, Edwards YJ, Han MS, Cavanagh-Kyros J, Barrett T, Kim JK, Davis RJ (2016) An alternative splicing program promotes adipose tissue thermogenesis. ELife, 5, e17672.

Wang SY, Cheng YY, Liu SC, Xu YX, Gao Y, Wang CL, Wang ZG, Feng TQ, Lu GH, Song J, Xia PJ, Hao LL (2021) A synonymous mutation in IGF-1 impacts the transcription and translation process of gene expression. Molecular Therapy - Nucleic Acids, 26, 1446– 1465.

Wang Y, Liu J, Huang B, Xu YM, Li J, Huang LF et al. (2015) Mechanism of alternative splicing and its regulation (Review). Biomedical Reports, 3(2), 152–158.

Wang Z, Burge CB (2008) Splicing regulation: From a parts list of regulatory elements to an integrated splicing code. RNA, 14(5), 802–813.

White RJ, Mackay E, Wilson SW, Busch-Nentwich EM (2022) Allele-specific gene expression can underlie altered transcript abundance in zebrafish mutants. ELife, 11, e72825.

Wittkopp PJ, Haerum BK, Clark AG (2004) Evolutionary changes in cis and trans gene regulation. Nature, 430(6995), 85–88.

Xu J (2004) Genotype-Environment Interactions of Spontaneous Mutations for Vegetative Fitness in the Human Pathogenic Fungus *Cryptococcus neoformans*. Genetics, 168(3), 1177–1188.

Xu S, Schaack S, Seyfert A, Choi E, Lynch M, Cristescu ME (2012) High Mutation Rates in the Mitochondrial Genomes of *Daphnia pulex*. Molecular Biology and Evolution, 29(2), 763–769.

Zalts H, Yanai I (2017) Developmental constraints shape the evolution of the nematode mid-developmental transition. Nature Ecology & Evolution, 1(5), Article 5.

