## Supplemental Material for "The effects of mutations on gene expression and alternative splicing: a case study of EMS-induced heritable mutations in the microcrustacean *Daphnia*"

**SUPPLEMENTARY MATERIAL**

**Supplementary Table S1**. Summary of whole-genome DNA sequencing data for each mutant line with, mapping rate and sequencing depth.

| **Mutant line** | **Total Sequences** | **Mapped reads** | **Mapping rate (%)** | **Average Depth** |
| --- | --- | --- | --- | --- |
| P3-1 | 27876025 | 24108745 | 86.49 | 20.7 |
| P3-2 | 39453960 | 28728608 | 72.82 | 24.4 |
| PP-1 | 38439774 | 33372636 | 86.82 | 28.5 |
| PP-2 | 38320557 | 33296967 | 86.89 | 28.3 |
| RB-1 | 38401662 | 31664358 | 82.46 | 26.9 |
| RB-2 | 33778759 | 28540895 | 84.49 | 24.3 |
| RY-1 | 39329907 | 31304410 | 79.59 | 26.7 |
| RY-2 | 38492505 | 32728183 | 85.02 | 28.1 |
| G-1 | 31372511 | 27797569 | 88.6 | 23.4 |
| G-2 | 32425011 | 27524999 | 84.89 | 22.8 |
| P-1 | 33458597 | 28904143 | 86.39 | 24.4 |
| PY3-1 | 31732028 | 27582657 | 86.92 | 23.2 |
| PY3-2 | 32518996 | 27936904 | 85.91 | 23.5 |
| R-1 | 31786328 | 28539697 | 89.79 | 24.0 |
| R-2 | 31278659 | 28378273 | 90.73 | 24.1 |
| Y-1 | 32457899 | 29329067 | 90.36 | 24.6 |
| **Average** | **34445199** | **29358632** | **85.51** | **24.9** |

**Supplementary Table S2**. Summary of RNA sequencing data for mutant lines.

| **Mutant**  **line** | **Total sequences** | **After trimming** | **% Retained** | **Total Alignments** | **Mapping rate** |
| --- | --- | --- | --- | --- | --- |
| BR-1-1 | 14479195 | 14478436 | 99.995 | 12951968 | 89.457 |
| BR-1-2 | 16932167 | 16930930 | 99.993 | 16192746 | 95.640 |
| BR-1-3 | 14189028 | 14188059 | 99.993 | 13275258 | 93.566 |
| BR-2-1 | 16220129 | 16219008 | 99.993 | 15434890 | 95.165 |
| BR-2-2 | 19217575 | 19216143 | 99.993 | 18366658 | 95.579 |
| BR-2-3 | 17846654 | 17845552 | 99.994 | 16977986 | 95.138 |
| G-1-1 | 24459488 | 24455295 | 99.983 | 23487807 | 96.044 |
| G-1-2 | 30819608 | 30812005 | 99.975 | 29367458 | 95.312 |
| G-1-3 | 30651687 | 30635229 | 99.946 | 28050037 | 91.561 |
| G-2-1 | 36943663 | 36911220 | 99.912 | 33825831 | 91.641 |
| G-2-2 | 24740613 | 24706683 | 99.863 | 21476386 | 86.925 |
| G-2-3 | 24025269 | 23963266 | 99.742 | 22500249 | 93.895 |
| P-1-1 | 21769002 | 21750579 | 99.915 | 20632572 | 94.860 |
| P-1-2 | 20165358 | 20158104 | 99.964 | 19121744 | 94.859 |
| P-1-3 | 20324587 | 20318905 | 99.972 | 19314091 | 95.055 |
| P3-1-1 | 12237329 | 12236352 | 99.992 | 11791275 | 96.363 |
| P3-1-2 | 15468266 | 15467375 | 99.994 | 14837156 | 95.925 |
| P3-1-3 | 22413807 | 22411779 | 99.991 | 21086238 | 94.086 |
| P3-2-1 | 18298659 | 18297673 | 99.995 | 17773108 | 97.133 |
| P3-2-2 | 19651101 | 19649799 | 99.993 | 19064869 | 97.023 |
| P3-2-3 | 21319219 | 21317714 | 99.993 | 20597702 | 96.622 |
| PP-1-1 | 14945627 | 14944124 | 99.990 | 13438416 | 89.924 |
| PP-1-2 | 15480770 | 15479392 | 99.991 | 14803742 | 95.635 |
| PP-1-3 | 18436550 | 18434983 | 99.992 | 17471538 | 94.774 |
| PP-2-1 | 22502559 | 22495319 | 99.968 | 20866653 | 92.760 |
| PP-2-2 | 18596333 | 18594022 | 99.988 | 17746164 | 95.440 |
| PP-2-3 | 26090199 | 26084679 | 99.979 | 23932493 | 91.749 |
| R-1-1 | 20824328 | 20815102 | 99.956 | 20232898 | 97.203 |
| R-1-2 | 22603348 | 22594946 | 99.963 | 22100482 | 97.812 |
| R-1-3 | 24283518 | 24271625 | 99.951 | 24520442 | 101.022 |
| R-2-1 | 23931170 | 23926138 | 99.979 | 23176052 | 96.865 |
| R-2-2 | 19131280 | 19123768 | 99.961 | 18298671 | 95.685 |
| R-2-3 | 22231762 | 22215201 | 99.926 | 21068811 | 94.840 |
| Y-1-1 | 18655231 | 18649148 | 99.967 | 17866136 | 95.801 |
| Y-1-2 | 15526481 | 15523321 | 99.980 | 14936343 | 96.219 |
| Y-1-3 | 15725993 | 15720326 | 99.964 | 15134673 | 96.275 |
| PY3-1-1 | 16353705 | 16350553 | 99.981 | 15576268 | 95.264 |
| PY3-1-2 | 21144915 | 21141396 | 99.983 | 19886973 | 94.067 |
| PY3-1-3 | 16455933 | 16452396 | 99.979 | 15633173 | 95.021 |
| PY3-2-1 | 20896695 | 20891799 | 99.977 | 19905749 | 95.280 |
| PY3-2-2 | 17672154 | 17668746 | 99.981 | 16841622 | 95.319 |
| PY3-2-3 | 16284381 | 16281382 | 99.982 | 15526272 | 95.362 |
| RY-1-1 | 17819218 | 17818332 | 99.995 | 16296216 | 91.458 |
| RY-1-2 | 24091306 | 24090139 | 99.995 | 22043782 | 91.505 |
| RY-1-3 | 14803027 | 14802171 | 99.994 | 13461606 | 90.943 |
| RY-2-1 | 22069817 | 22068348 | 99.993 | 20957309 | 94.965 |
| RY-2-2 | 26069363 | 26067705 | 99.994 | 24788003 | 95.091 |
| RY-2-3 | 15774528 | 15773551 | 99.994 | 14943889 | 94.740 |
| AR-1 | 25962592 | 25956036 | 99.975 | 24346962 | 93.801 |
| AR-2 | 12452945 | 12449443 | 99.972 | 12001705 | 96.404 |
| AR-3 | 19289167 | 19285232 | 99.980 | 18548829 | 96.182 |
| **Average** | **20162300** | **20155675.08** | **99.97** | **19068194.14** | **94.69** |

**Supplementary Table S3.** Summary statistics of the EMS-induced heritable mutations. All mutation rates are for one generation.

| **Mutant line** | **No. of EMS-induced mutations** | **Mutation rate** | **No. of mutations within coding regions** | **Per gene mutation rate** | **No. of non-synonymous mutations** | **Non-synonymous mutation rate** | **No. of synonymous mutations** | **Ratio of non-synonymous/**  **synonymous**  **mutation** |
| --- | --- | --- | --- | --- | --- | --- | --- | --- |
| P3-1 | 133 | 1.61×10^-6^ | 69 | 0.0029 | 21 | 0.00084 | 22 | 0.95 |
| P3-2 | 146 | 1.77×10^-6^ | 80 | 0.0033 | 26 | 0.00104 | 21 | 1.24 |
| PP-1 | 132 | 1.60×10^-6^ | 52 | 0.0022 | 24 | 0.00096 | 12 | 2 |
| PP-2 | 130 | 1.57×10^-6^ | 49 | 0.002 | 24 | 0.00096 | 12 | 2 |
| RB-1 | 131 | 1.59×10^-6^ | 72 | 0.003 | 31 | 0.00124 | 15 | 2.07 |
| RB-2 | 125 | 1.51×10^-6^ | 69 | 0.0029 | 30 | 0.0012 | 13 | 2.31 |
| RY-1 | 570 | 6.90×10^-6^ | 319 | 0.013 | 140 | 0.0056 | 72 | 1.94 |
| RY-2 | 576 | 6.98×10^-6^ | 320 | 0.013 | 114 | 0.00456 | 68 | 1.68 |
| G-1 | 190 | 2.30×10^-6^ | 91 | 0.0038 | 34 | 0.00136 | 18 | 1.89 |
| G-2 | 182 | 2.20×10^-6^ | 84 | 0.0035 | 28 | 0.00112 | 19 | 1.47 |
| P-1 | 169 | 2.05×10^-6^ | 87 | 0.0036 | 40 | 0.0016 | 21 | 1.90 |
| PY3-1 | 130 | 1.57×10^-6^ | 63 | 0.0026 | 27 | 0.00108 | 6 | 4.5 |
| PY3-2 | 130 | 1.57×10^-6^ | 60 | 0.0025 | 22 | 0.00088 | 7 | 3.14 |
| R-1 | 158 | 1.91×10^-6^ | 72 | 0.003 | 29 | 0.00116 | 12 | 2.42 |
| R-2 | 167 | 2.02×10^-6^ | 75 | 0.0031 | 31 | 0.00124 | 14 | 2.21 |
| Y-1 | 181 | 2.19×10^-6^ | 89 | 0.0037 | 25 | 0.001 | 18 | 1.39 |

**Supplementary Table S4.** The number of different types of base substitutions per mutant line.

| **Base**  **Substitution** | **G-1** | **G-2** | **P3-1** | **P3-2** | **PP-1** | **PP-2** | **P-1** | **PY3-1** | **PY3-2** | **RB-1** | **RB-2** | **R-1** | **R-2** | **RY-1** | **RY-2** | **Y-1** |
| --- | --- | --- | --- | --- | --- | --- | --- | --- | --- | --- | --- | --- | --- | --- | --- | --- |
| A>C: | 1 | 1 | 0 | 0 | 1 | 1 | 0 | 0 | 0 | 0 | 0 | 0 | 0 | 0 | 1 | 0 |
| A>G: | 1 | 1 | 2 | 1 | 0 | 0 | 0 | 1 | 1 | 1 | 1 | 1 | 2 | 3 | 2 | 0 |
| A>T: | 4 | 6 | 1 | 1 | 0 | 0 | 0 | 0 | 0 | 2 | 2 | 0 | 0 | 9 | 10 | 1 |
| C>A: | 5 | 6 | 5 | 6 | 3 | 4 | 5 | 1 | 2 | 4 | 5 | 2 | 2 | 8 | 6 | 5 |
| C>G: | 3 | 3 | 3 | 3 | 2 | 3 | 0 | 1 | 3 | 0 | 0 | 1 | 1 | 0 | 0 | 1 |
| C>T: | 83 | 80 | 68 | 78 | 49 | 48 | 94 | 51 | 53 | 61 | 60 | 65 | 66 | 282 | 287 | 87 |
| G>A: | 81 | 82 | 65 | 63 | 74 | 71 | 76 | 79 | 75 | 55 | 55 | 94 | 100 | 301 | 304 | 89 |
| G>C: | 0 | 0 | 1 | 1 | 0 | 0 | 0 | 1 | 0 | 0 | 0 | 0 | 0 | 1 | 1 | 2 |
| G>T: | 9 | 7 | 4 | 7 | 4 | 4 | 6 | 4 | 4 | 12 | 10 | 5 | 4 | 9 | 8 | 5 |
| T>A: | 8 | 5 | 0 | 1 | 1 | 2 | 2 | 0 | 0 | 0 | 0 | 1 | 1 | 9 | 11 | 4 |
| T>C: | 2 | 2 | 0 | 0 | 2 | 1 | 0 | 2 | 2 | 0 | 0 | 1 | 1 | 4 | 6 | 3 |
| T>G: | 3 | 1 | 0 | 1 | 0 | 0 | 2 | 0 | 0 | 0 | 0 | 0 | 0 | 0 | 0 | 0 |
| **Ts/Tv ratio:** | **5.1** | **5.7** | **9.6** | **7.1** | **11.4** | **8.6** | **11.3** | **19** | **14.6** | **6.5** | **6.8** | **17.9** | **21.1** | **16.4** | **16.2** | **9.9** |
| **Total**  **mutations:** | **190** | **182** | **133** | **146** | **132** | **130** | **169** | **130** | **130** | **131** | **125** | **158** | **167** | **570** | **576** | **181** |

**Supplementary Table S5:** Summary statistics of the differentially expressed genes and DE genes containing mutations. The Benjamini-Hochberg method (FDR < 0.05) was used to adjust the p-values for multiple testing, and genes with a p-value <0.05 were considered significantly differentially expressed.

| **Mutant**  **line** | **DE genes** | **Upregulated** | **Downregulated** | **Ratio of down-/ upregulated genes** | **DE genes with mutations** | **Upregulated genes with mutations** | **Downregulated genes with mutations** | **Ratio of down-/ upregulated genes with mutations** |
| --- | --- | --- | --- | --- | --- | --- | --- | --- |
| P3-1 | 4990 | 2349 | 2641 | 1.12 | 76 | 40 | 36 | 0.90 |
| P3-2 | 6280 | 2971 | 3309 | 1.11 | 100 | 59 | 41 | 0.69 |
| PP-1 | 1176 | 254 | 922 | 3.63 | 5 | 0 | 5 | - |
| PP-2 | 3562 | 1412 | 2150 | 1.52 | 37 | 15 | 22 | 1.47 |
| RB-1 | 3484 | 1751 | 1733 | 0.99 | 53 | 35 | 18 | 0.51 |
| RB-2 | 6470 | 3109 | 3361 | 1.08 | 87 | 48 | 39 | 0.81 |
| RY-1 | 6444 | 3143 | 3301 | 1.05 | 393 | 214 | 179 | 0.84 |
| RY-2 | 6606 | 3133 | 3473 | 1.11 | 423 | 236 | 187 | 0.79 |
| G-1 | 2347 | 980 | 1367 | 1.39 | 36 | 15 | 21 | 1.4 |
| G-2 | 4504 | 2034 | 2470 | 1.21 | 76 | 32 | 44 | 1.38 |
| P-1 | 3527 | 1680 | 1847 | 1.10 | 48 | 32 | 16 | 0.5 |
| PY3-1 | 1750 | 524 | 1226 | 2.34 | 24 | 6 | 18 | 3 |
| PY3-2 | 6084 | 2860 | 3224 | 1.13 | 74 | 38 | 36 | 0.94 |
| R-1 | 3046 | 1085 | 1961 | 1.81 | 44 | 18 | 26 | 1.44 |
| R-2 | 2830 | 1159 | 1671 | 1.44 | 43 | 19 | 24 | 1.26 |
| Y-1 | 1937 | 705 | 1232 | 1.75 | 37 | 15 | 22 | 1.47 |

**Supplementary Table S6**. The number of mutations found in different genomic regions across all mutant lines.

| **Effect**  **Region** | **PP-1** | **PP-2** | **P3-1** | **P3-2** | **RB-1** | **RB-2** | **RY-1** | **RY-2** | **G-1** | **G-2** | **P-1** | **PY3-1** | **PY3-2** | **R-1** | **R-2** | **Y-1** | **Total** |
| --- | --- | --- | --- | --- | --- | --- | --- | --- | --- | --- | --- | --- | --- | --- | --- | --- | --- |
| Downstream | 116 | 122 | 123 | 128 | 119 | 116 | 536 | 543 | 192 | 176 | 168 | 130 | 131 | 134 | 151 | 162 | 3047 |
| Exon | 36 | 36 | 41 | 45 | 44 | 41 | 224 | 225 | 55 | 50 | 61 | 39 | 33 | 42 | 46 | 50 | 1068 |
| Intergenic | 83 | 83 | 71 | 74 | 67 | 66 | 281 | 289 | 104 | 107 | 92 | 75 | 78 | 91 | 95 | 106 | 1762 |
| Intron | 17 | 13 | 25 | 30 | 19 | 19 | 98 | 97 | 40 | 32 | 26 | 21 | 24 | 27 | 28 | 22 | 538 |
| Splice  site region | 1 | 1 | 7 | 7 | 5 | 5 | 8 | 8 | 3 | 2 | 1 | 2 | 0 | 1 | 4 | 3 | 58 |
| Upstream | 123 | 124 | 103 | 117 | 126 | 124 | 589 | 593 | 186 | 193 | 171 | 120 | 121 | 136 | 151 | 165 | 3142 |
| UTR 3 prime | 8 | 7 | 8 | 12 | 3 | 3 | 23 | 24 | 6 | 4 | 7 | 3 | 3 | 10 | 9 | 14 | 144 |
| UTR 5 prime | 4 | 4 | 4 | 4 | 9 | 9 | 14 | 13 | 10 | 10 | 7 | 6 | 9 | 15 | 15 | 14 | 147 |

**Supplementary Table S7.** The number of mutations found in different genomic regions for differentially expressed (DE) genes.

| **Effect Region** | **PP-1** | **PP-2** | **P3-1** | **P3-2** | **RB-1** | **RB-2** | **RY-1** | **RY-2** | **G-1** | **G-2** | **P-1** | **PY3-1** | **PY3-2** | **R-1** | **R-2** | **Y-1** | **Total** |
| --- | --- | --- | --- | --- | --- | --- | --- | --- | --- | --- | --- | --- | --- | --- | --- | --- | --- |
| Downstream | 1 | 21 | 28 | 33 | 31 | 41 | 160 | 169 | 14 | 22 | 24 | 15 | 36 | 14 | 16 | 25 | 650 |
| Exon | 0 | 8 | 19 | 24 | 8 | 14 | 83 | 87 | 6 | 15 | 11 | 3 | 11 | 10 | 8 | 4 | 311 |
| Intron | 0 | 3 | 11 | 15 | 3 | 8 | 39 | 38 | 1 | 7 | 5 | 0 | 5 | 2 | 2 | 1 | 140 |
| Splice site region | 0 | 0 | 3 | 4 | 0 | 0 | 3 | 2 | 0 | 0 | 0 | 0 | 0 | 1 | 3 | 0 | 16 |
| Upstream | 4 | 19 | 27 | 47 | 18 | 29 | 176 | 192 | 23 | 42 | 19 | 9 | 29 | 24 | 28 | 12 | 698 |
| UTR 3 prime | 0 | 1 | 1 | 2 | 0 | 0 | 6 | 7 | 2 | 3 | 4 | 0 | 1 | 7 | 7 | 2 | 43 |
| UTR 5 prime | 0 | 2 | 3 | 3 | 0 | 4 | 4 | 2 | 0 | 2 | 0 | 1 | 2 | 3 | 3 | 4 | 33 |

**Supplementary Table S8.** Number of mutations found in different genomic regions for non-differentially expressed (non-DE) genes.

| **Effect Region** | **PP-1** | **PP-2** | **P3-1** | **P3-2** | **RB-1** | **RB-2** | **RY-1** | **RY-2** | **G-1** | **G-2** | **P-1** | **PY3-1** | **PY3-2** | **R-1** | **R-2** | **Y-1** | **Total** |
| --- | --- | --- | --- | --- | --- | --- | --- | --- | --- | --- | --- | --- | --- | --- | --- | --- | --- |
| Downstream | 115 | 101 | 95 | 95 | 88 | 75 | 376 | 374 | 178 | 174 | 144 | 115 | 95 | 120 | 135 | 137 | 2397 |
| Exon | 36 | 28 | 22 | 21 | 36 | 27 | 141 | 138 | 49 | 35 | 50 | 36 | 22 | 32 | 38 | 46 | 757 |
| Intron | 17 | 10 | 14 | 15 | 16 | 11 | 59 | 59 | 39 | 25 | 21 | 21 | 19 | 25 | 26 | 21 | 398 |
| Splice site region | 1 | 1 | 4 | 3 | 5 | 5 | 5 | 6 | 3 | 2 | 1 | 2 | 0 | 1 | 1 | 3 | 42 |
| Upstream | 119 | 105 | 76 | 70 | 108 | 95 | 413 | 401 | 163 | 151 | 152 | 111 | 92 | 112 | 123 | 153 | 2444 |
| UTR 3 prime | 8 | 6 | 7 | 10 | 3 | 3 | 17 | 17 | 4 | 1 | 3 | 3 | 2 | 3 | 2 | 11 | 101 |
| UTR 5 prime | 4 | 2 | 1 | 1 | 9 | 5 | 10 | 11 | 10 | 8 | 7 | 5 | 7 | 12 | 12 | 10 | 114 |

**Supplementary Table S9.** Impact of mutations located in differentially expressed genes.

| **Mutant line** | **Direction of regulation** | **High** | **Low** | **Moderate** | **Modifier** |
| --- | --- | --- | --- | --- | --- |
| G-1 | Upregulation | 0 | 0 | 1 | 18 |
|  | Downregulation | 0 | 3 | 2 | 22 |
| G-2 | Upregulation | 0 | 3 | 3 | 30 |
|  | Downregulation | 2 | 1 | 6 | 46 |
| P3-1 | Upregulation | 2 | 7 | 3 | 37 |
|  | Downregulation | 0 | 4 | 4 | 33 |
| P3-2 | Upregulation | 2 | 12 | 9 | 53 |
|  | Downregulation | 0 | 2 | 1 | 46 |
| PP-1 | Downregulation | 0 | 0 | 0 | 5 |
| PP-2 | Upregulation | 0 | 2 | 1 | 21 |
|  | Downregulation | 0 | 1 | 4 | 25 |
| P-1 | Upregulation | 0 | 5 | 5 | 33 |
|  | Downregulation | 0 | 0 | 1 | 19 |
| PY3-1 | Upregulation | 0 | 0 | 0 | 9 |
|  | Downregulation | 0 | 1 | 2 | 16 |
| PY3-2 | Upregulation | 2 | 3 | 5 | 35 |
|  | Downregulation | 0 | 0 | 1 | 38 |
| RB-1 | Upregulation | 0 | 1 | 5 | 35 |
|  | Downregulation | 0 | 0 | 2 | 17 |
| RB-2 | Upregulation | 0 | 2 | 7 | 44 |
|  | Downregulation | 0 | 3 | 3 | 37 |
| R-1 | Upregulation | 0 | 0 | 5 | 23 |
|  | Downregulation | 1 | 3 | 2 | 26 |
| R-2 | Upregulation | 0 | 0 | 2 | 29 |
|  | Downregulation | 1 | 2 | 4 | 26 |
| RY-1 | Upregulation | 5 | 25 | 37 | 184 |
|  | Downregulation | 1 | 7 | 12 | 197 |
| RY-2 | Upregulation | 7 | 21 | 45 | 197 |
|  | Downregulation | 1 | 5 | 11 | 208 |
| Y-1 | Upregulation | 3 | 0 | 1 | 16 |
|  | Downregulation | 0 | 0 | 0 | 28 |
| **Total** |  | **27** | **113** | **184** | **1553** |

**Supplementary Table S10**. Effect of mutations impacting differentially expressed genes.

| **Mutant**  **line** | **Direction**  **of**  **Regulation** | **3’ UTR** | **5’ UTR premature start codon gain** | **5’ UTR** | **Down-stream** | **Intron** | **Mis-sense** | **Start-lost** | **Splice acceptor** | **Splice region** | **Stop gained** | **Synonymous** | **Up-stream** |
| --- | --- | --- | --- | --- | --- | --- | --- | --- | --- | --- | --- | --- | --- |
| G-1 | Up | 0 | 0 | 0 | 4 | 0 | 1 | 0 | 0 | 0 | 0 | 0 | 14 |
|  | Down | 2 | 0 | 0 | 10 | 1 | 2 | 0 | 0 | 0 | 0 | 3 | 9 |
| G-2 | Up | 2 | 0 | 1 | 9 | 1 | 3 | 0 | 0 | 0 | 0 | 3 | 17 |
|  | Down | 1 | 0 | 1 | 13 | 6 | 6 | 0 | 0 | 0 | 2 | 1 | 25 |
| P3-1 | Up | 1 | 0 | 3 | 13 | 6 | 3 | 1 | 0 | 2 | 1 | 6 | 14 |
|  | Down | 0 | 0 | 0 | 15 | 5 | 4 | 0 | 0 | 1 | 0 | 4 | 13 |
| P3-2 | Up | 2 | 0 | 3 | 18 | 8 | 9 | 1 | 0 | 3 | 1 | 10 | 23 |
|  | Down | 0 | 0 | 0 | 15 | 7 | 1 | 0 | 0 | 1 | 0 | 2 | 24 |
| PP-1 | Down | 0 | 0 | 0 | 1 | 0 | 0 | 0 | 0 | 0 | 0 | 0 | 4 |
| PP-2 | Up | 0 | 0 | 0 | 10 | 0 | 1 | 0 | 0 | 0 | 0 | 2 | 11 |
|  | Down | 1 | 0 | 2 | 11 | 3 | 4 | 0 | 0 | 0 | 0 | 1 | 8 |
| P-1 | Up | 4 | 0 | 0 | 17 | 4 | 5 | 0 | 0 | 0 | 0 | 5 | 8 |
|  | Down | 0 | 0 | 0 | 7 | 1 | 1 | 0 | 0 | 0 | 0 | 0 | 11 |
| PY3-1 | Up | 0 | 0 | 0 | 6 | 0 | 0 | 0 | 0 | 0 | 0 | 0 | 3 |
|  | Down | 0 | 0 | 1 | 9 | 0 | 2 | 0 | 0 | 0 | 0 | 1 | 6 |
| PY3-2 | Up | 1 | 0 | 1 | 16 | 4 | 5 | 0 | 0 | 0 | 2 | 3 | 13 |
|  | Down | 0 | 0 | 1 | 20 | 1 | 1 | 0 | 0 | 0 | 0 | 0 | 16 |
| RB-1 | Up | 0 | 0 | 0 | 22 | 1 | 5 | 0 | 0 | 0 | 0 | 1 | 12 |
|  | Down | 0 | 0 | 0 | 9 | 2 | 2 | 0 | 0 | 0 | 0 | 0 | 6 |
| RB-2 | Up | 0 | 0 | 2 | 19 | 7 | 7 | 0 | 0 | 0 | 0 | 2 | 16 |
|  | Down | 0 | 1 | 1 | 22 | 1 | 3 | 0 | 0 | 0 | 0 | 2 | 13 |
| R-1 | Up | 6 | 0 | 1 | 4 | 1 | 5 | 0 | 0 | 0 | 0 | 0 | 11 |
|  | Down | 1 | 0 | 2 | 10 | 1 | 2 | 0 | 0 | 1 | 1 | 2 | 13 |
| R-2 | Up | 6 | 0 | 0 | 9 | 0 | 2 | 0 | 0 | 0 | 0 | 0 | 14 |
|  | Down | 1 | 0 | 3 | 7 | 2 | 4 | 0 | 0 | 3 | 1 | 1 | 14 |
| RY-1 | Up | 4 | 1 | 3 | 70 | 31 | 37 | 0 | 0 | 2 | 5 | 22 | 78 |
|  | Down | 2 | 0 | 0 | 90 | 8 | 12 | 0 | 1 | 0 | 0 | 7 | 98 |
| RY-2 | Up | 5 | 1 | 1 | 73 | 29 | 45 | 0 | 0 | 1 | 7 | 19 | 90 |
|  | Down | 2 | 0 | 0 | 96 | 9 | 11 | 0 | 1 | 0 | 0 | 5 | 102 |
| Y-1 | Up | 2 | 0 | 2 | 5 | 1 | 1 | 0 | 0 | 0 | 3 | 0 | 6 |
|  | Down | 0 | 0 | 2 | 20 | 0 | 0 | 0 | 0 | 0 | 0 | 0 | 6 |
| **Total** |  | **43** | **3** | **30** | **650** | **140** | **184** | **2** | **2** | **14** | **23** | **102** | **698** |

**Supplementary Table S11.** Impact of mutations located in non-differentially expressed (non-DE) genes.

| **Mutant line** | **High** | **Low** | **Moderate** | **Modifier** |
| --- | --- | --- | --- | --- |
| **G-1** | 3 | 18 | 31 | 394 |
| **G-2** | 1 | 17 | 19 | 339 |
| **P3-1** | 0 | 16 | 14 | 193 |
| **P3-2** | 0 | 12 | 16 | 191 |
| **PP-1** | 0 | 14 | 24 | 262 |
| **PP-2** | 0 | 11 | 19 | 223 |
| **P-1** | 0 | 17 | 34 | 327 |
| **PY3-1** | 6 | 9 | 25 | 253 |
| **PY3-2** | 2 | 7 | 16 | 212 |
| **RB-1** | 1 | 20 | 24 | 223 |
| **RB-2** | 1 | 14 | 20 | 189 |
| **R-1** | 0 | 12 | 22 | 270 |
| **R-2** | 0 | 15 | 25 | 297 |
| **RY-1** | 7 | 48 | 91 | 875 |
| **RY-2** | 6 | 51 | 58 | 861 |
| **Y-1** | 5 | 22 | 24 | 332 |
| **Total** | **32** | **303** | **462** | **5441** |

**Supplementary Table S12**. Effect of mutations impacting non-differentially expressed (non-DE) genes.

| **Mutant**  **line** | **3’ UTR** | **5’ UTR premature start codon gain** | **5’ UTR** | **Down-stream** | **Intron** | **Mis-sense** | **Start-lost** | **Splice region** | **Stop gained** | **Synonymous** | **Up-stream** |
| --- | --- | --- | --- | --- | --- | --- | --- | --- | --- | --- | --- |
| G-1 | 4 | 0 | 10 | 178 | 39 | 31 | 0 | 3 | 3 | 15 | 163 |
| G-2 | 1 | 0 | 8 | 154 | 25 | 19 | 0 | 2 | 1 | 15 | 151 |
| P3-1 | 7 | 0 | 1 | 95 | 14 | 14 | 0 | 4 | 0 | 12 | 76 |
| P3-2 | 10 | 0 | 1 | 95 | 15 | 16 | 0 | 3 | 0 | 9 | 70 |
| PP-1 | 8 | 1 | 3 | 115 | 17 | 24 | 0 | 1 | 0 | 12 | 119 |
| PP-2 | 6 | 1 | 1 | 101 | 10 | 19 | 0 | 1 | 0 | 9 | 105 |
| P-1 | 3 | 0 | 7 | 144 | 21 | 34 | 0 | 1 | 0 | 16 | 152 |
| PY3-1 | 3 | 2 | 3 | 115 | 21 | 25 | 2 | 2 | 4 | 5 | 111 |
| PY3-2 | 2 | 3 | 4 | 95 | 19 | 16 | 1 | 0 | 1 | 4 | 92 |
| RB-1 | 3 | 1 | 8 | 88 | 16 | 24 | 0 | 5 | 1 | 14 | 108 |
| RB-2 | 3 | 0 | 5 | 75 | 11 | 20 | 0 | 5 | 1 | 9 | 95 |
| R-1 | 3 | 2 | 10 | 120 | 25 | 22 | 0 | 0 | 0 | 10 | 112 |
| R-2 | 2 | 1 | 11 | 135 | 26 | 25 | 0 | 1 | 0 | 13 | 123 |
| RY-1 | 17 | 0 | 10 | 376 | 59 | 91 | 0 | 5 | 7 | 43 | 413 |
| RY-2 | 17 | 1 | 10 | 374 | 59 | 58 | 0 | 6 | 6 | 44 | 401 |
| Y-1 | 12 | 1 | 9 | 137 | 21 | 24 | 0 | 3 | 5 | 18 | 153 |
| **Total** | **101** | **13** | **101** | **2397** | **398** | **462** | **3** | **42** | **29** | **248** | **2444** |

**Supplementary Table S13**. Alternatively spliced events detected across all 16 EMS mutant lines.

| **Mutant line** | **No. of AS genes** | **A3SS** | **A5SS** | **MXE** | **RI** | **SE** |
| --- | --- | --- | --- | --- | --- | --- |
| P3-1 | 381 | 45 | 31 | 88 | 99 | 118 |
| P3-2 | 415 | 37 | 33 | 82 | 104 | 159 |
| PP-1 | 282 | 26 | 39 | 52 | 83 | 82 |
| PP-2 | 363 | 34 | 45 | 66 | 100 | 118 |
| RB-1 | 212 | 17 | 16 | 30 | 69 | 80 |
| RB-2 | 404 | 45 | 48 | 78 | 126 | 107 |
| RY-1 | 522 | 48 | 58 | 83 | 175 | 158 |
| RY-2 | 593 | 55 | 59 | 98 | 166 | 215 |
| G-1 | 415 | 38 | 38 | 59 | 188 | 92 |
| G-2 | 627 | 54 | 69 | 73 | 250 | 181 |
| P-1 | 419 | 50 | 33 | 50 | 180 | 106 |
| PY3-1 | 241 | 36 | 24 | 45 | 71 | 65 |
| PY3-2 | 436 | 55 | 61 | 59 | 130 | 131 |
| R-1 | 334 | 46 | 31 | 71 | 84 | 102 |
| R-2 | 349 | 43 | 38 | 62 | 123 | 83 |
| Y-1 | 295 | 36 | 42 | 55 | 79 | 83 |
| **Total** | **6288** | **665** | **665** | **1051** | **2027** | **1880** |

**Supplementary Table S14**. Alternatively spliced events of genes containing an EMS-induced mutation in each mutant line.

| **Mutant line** | **Total** | **A3SS** | **A5SS** | **MXE** | **RI** | **SE** | **AS genes also DE** |
| --- | --- | --- | --- | --- | --- | --- | --- |
| P3-1 | 17 | 3 | 0 | 3 | 0 | 11 | 3 |
| P3-2 | 8 | 0 | 0 | 0 | 0 | 8 | 1 |
| PP-1 | 11 | 1 | 0 | 4 | 2 | 4 | 0 |
| PP-2 | 2 | 0 | 0 | 0 | 2 | 0 | 2 |
| RB-1 | 10 | 0 | 1 | 3 | 2 | 4 | 4 |
| RB-2 | 23 | 2 | 2 | 12 | 2 | 5 | 4 |
| RY-1 | 42 | 5 | 7 | 1 | 11 | 18 | 11 |
| RY-2 | 57 | 5 | 6 | 4 | 10 | 32 | 12 |
| G-1 | 16 | 0 | 1 | 6 | 6 | 3 | 0 |
| G-2 | 16 | 0 | 2 | 5 | 3 | 6 | 2 |
| P-1 | 12 | 4 | 0 | 0 | 7 | 1 | 2 |
| PY3-1 | 3 | 0 | 1 | 0 | 0 | 2 | 1 |
| PY3-2 | 4 | 0 | 2 | 0 | 2 | 0 | 1 |
| R-1 | 10 | 0 | 0 | 4 | 4 | 2 | 0 |
| R-2 | 15 | 0 | 0 | 6 | 3 | 6 | 1 |
| Y-1 | 3 | 0 | 0 | 0 | 1 | 2 | 0 |
| **Total** | **249** | **20** | **22** | **48** | **55** | **104** | **44** |

**Supplementary Table S15**. Alternatively spliced events of genes without an EMS-induced mutation in each mutant line.

| **Mutant line** | **Total** | **A3SS** | **A5SS** | **MXE** | **RI** | **SE** |
| --- | --- | --- | --- | --- | --- | --- |
| P3-1 | 364 | 42 | 31 | 85 | 99 | 107 |
| P3-2 | 407 | 37 | 33 | 82 | 104 | 151 |
| PP-1 | 271 | 25 | 39 | 48 | 81 | 78 |
| PP-2 | 361 | 34 | 45 | 66 | 98 | 118 |
| RB-1 | 202 | 17 | 15 | 27 | 67 | 76 |
| RB-2 | 381 | 43 | 46 | 66 | 124 | 102 |
| RY-1 | 480 | 43 | 51 | 82 | 164 | 140 |
| RY-2 | 536 | 50 | 53 | 94 | 156 | 183 |
| G-1 | 399 | 38 | 37 | 53 | 182 | 89 |
| G-2 | 611 | 54 | 67 | 68 | 247 | 175 |
| P-1 | 407 | 46 | 33 | 50 | 173 | 105 |
| PY3-1 | 238 | 36 | 23 | 45 | 71 | 63 |
| PY3-2 | 432 | 55 | 59 | 59 | 128 | 131 |
| R-1 | 324 | 46 | 31 | 67 | 80 | 100 |
| R-2 | 334 | 43 | 38 | 56 | 120 | 77 |
| Y-1 | 292 | 36 | 42 | 55 | 78 | 81 |
| **Total** | **6039** | **645** | **643** | **1003** | **1972** | **1776** |

**Supplementary Table S16.** Impact of mutations affecting alternatively spliced genes per mutant line.

| **Mutant line** | **High Impact** | **Low impact** | **Moderate impact** | **Modifier** |
| --- | --- | --- | --- | --- |
| G_1 | 0 | 5 | 0 | 10 |
| G_2 | 0 | 1 | 0 | 12 |
| P3_1 | 0 | 1 | 0 | 6 |
| P3_2 | 0 | 0 | 1 | 3 |
| PP_1 | 0 | 0 | 5 | 6 |
| PP_2 | 0 | 0 | 0 | 7 |
| P_1 | 0 | 0 | 3 | 20 |
| PY3_1 | 2 | 0 | 0 | 3 |
| PY3_2 | 0 | 0 | 1 | 5 |
| BR_1 | 0 | 0 | 2 | 6 |
| BR_2 | 0 | 0 | 1 | 9 |
| R_1 | 0 | 0 | 0 | 7 |
| R_2 | 0 | 0 | 0 | 9 |
| RY_1 | 2 | 2 | 2 | 29 |
| RY_2 | 2 | 4 | 5 | 25 |
| Y_1 | 0 | 1 | 0 | 2 |
| **Total** | **6** | **14** | **20** | **159** |

**Supplementary Table S17**. The number of mutations in different genomic regions impacting alternatively spliced genes.

| **Mutant line** | **3’ UTR** | **Downstream** | **Intron** | **Missense** | **Splice region** | **Splice acceptor** | **Splice donor** | **Stop-gained** | **Synonymous variant** | **Upstream** |
| --- | --- | --- | --- | --- | --- | --- | --- | --- | --- | --- |
| G-1 | 0 | 2 | 2 | 0 | 0 | 0 | 0 | 0 | 5 | 6 |
| G-2 | 0 | 3 | 2 | 0 | 0 | 0 | 0 | 0 | 1 | 7 |
| P3-1 | 0 | 1 | 1 | 0 | 0 | 0 | 0 | 0 | 1 | 4 |
| P3-2 | 0 | 0 | 0 | 1 | 0 | 0 | 0 | 0 | 0 | 3 |
| PP-1 | 0 | 0 | 0 | 5 | 0 | 0 | 0 | 0 | 0 | 6 |
| PP-2 | 0 | 0 | 0 | 0 | 0 | 0 | 0 | 0 | 0 | 7 |
| P-1 | 0 | 5 | 1 | 3 | 0 | 0 | 0 | 0 | 0 | 11 |
| PY3-1 | 0 | 3 | 0 | 0 | 0 | 0 | 0 | 2 | 0 | 0 |
| PY3-2 | 0 | 5 | 0 | 1 | 0 | 0 | 0 | 0 | 0 | 0 |
| BR-1 | 0 | 1 | 1 | 2 | 0 | 0 | 0 | 0 | 0 | 4 |
| BR-2 | 0 | 1 | 1 | 1 | 0 | 0 | 0 | 0 | 0 | 7 |
| R-1 | 0 | 3 | 1 | 0 | 0 | 0 | 0 | 0 | 0 | 3 |
| R-2 | 0 | 1 | 2 | 0 | 0 | 0 | 0 | 0 | 0 | 6 |
| RY-1 | 0 | 7 | 6 | 2 | 1 | 1 | 1 | 0 | 1 | 19 |
| RY-2 | 1 | 7 | 5 | 5 | 0 | 0 | 1 | 1 | 4 | 13 |
| Y-1 | 0 | 1 | 0 | 0 | 0 | 0 | 0 | 0 | 1 | 1 |
| **Total** | **1** | **40** | **22** | **20** | **1** | **1** | **2** | **3** | **13** | **97** |
